# ConsHMM Atlas: conservation state annotations for major genomes and human genetic variation

**DOI:** 10.1101/2020.03.01.955443

**Authors:** Adriana Arneson, Brooke Felsheim, Jennifer Chien, Jason Ernst

## Abstract

ConsHMM is a method recently introduced to annotate genomes into conservation states, which are defined based on the combinatorial and spatial patterns of which species align to and match a reference genome in a multi-species DNA sequence alignment. Previously, ConsHMM was only applied to a single genome for one multi-species sequence alignment. Here we apply ConsHMM to produce 22 additional genome annotations covering human and seven other organisms for a variety of multi-species alignments. Additionally, we have extended ConsHMM to generate allele specific annotations, which we used to produce conservation state annotations for every possible single nucleotide mutation in the human genome. Finally, we provide a web interface to interactively visualize parameters and annotation enrichments for ConsHMM models. These annotations and visualizations comprise the ConsHMM Atlas, which we expect will be a valuable resource for analyzing a variety of major genomes and genetic variation.

## Introduction

We recently introduced the ConsHMM method(Arneson and Ernst 2019) to annotate reference genomes at single-nucleotide resolution into a number of different ‘conservation states’ based on the combinatorial and spatial patterns of which species have a nucleotide aligning to and/or matching the reference genome in a multi-species DNA sequence alignment. To do this ConsHMM uses a multivariate hidden Markov model (HMM), building off the ChromHMM approach for modeling epigenomic data(Ernst and Kellis 2012), without making any explicit phylogenetic modeling assumptions. Each nucleotide in the reference genome receives an annotation corresponding to the state of the HMM with the maximum posterior probability.

ConsHMM annotations are complementary to previous whole genome comparative genomic annotations, which have primarily focused on univariate scores or binary element calls of constraint(Cooper et al. 2005; Siepel et al. 2005; Garber et al. 2009; Pollard et al. 2010). We previously applied ConsHMM to annotate one reference genome, human hg19, based on a 100-way vertebrate alignment(Arneson and Ernst 2019). The conservation states had diverse and biologically meaningful enrichments for other genomic annotations, and were also able to isolate putative artifacts in the underlying multiple sequence alignment, which can confound some traditional constraint annotations.

Here we report applying ConsHMM to produce an additional 22 genome annotations for different reference genomes and based on different multi-species DNA sequence alignments. In addition to human, seven other organisms are represented in these additional genome annotations. We have also extended the ConsHMM software to produce allele specific annotations opposed to only position specific annotations based on the reference allele. We have applied this to produce annotations for each possible single-nucleotide mutation for every nucleotide in the human genome. To aid in the analysis of different ConsHMM models, we have created a web-interface for interactive visualization of model parameters and annotation enrichments. These new annotations of the human genome and variation as well as model organism genomes along with a new visualization tool comprise the ConsHMM Atlas (http://www.biolchem.ucla.edu/labs/ernst/ConsHMMAtlas), which we expect to be a valuable resource to community for analyzing various genomes and genetic variation.

## Materials and Methods

### Generating ConsHMM annotations for reference genomes

We used ConsHMM v1.0 as described in Arneson and Ernst (2019) to learn models parameters, to generate segmentation and annotations of reference genomes, and to compute the enrichments for external annotations. We used the same parameters except the number of states parameters. The number of states we used for each alignment depended on the number of species in the alignment. Specifically, if the alignment had more than 50 species, then the number of states was equivalent to the number of species in the alignment; if the alignment had between 25 and 49 species, then the number of states was set to 50; if the alignment had less than 25 species, then the number of states was set to 25. This set of rules allows for the number of states to be dependent on the number of species in the alignment, while also ensuring a sufficient, but not excessive, number of states for alignments with smaller number of species.

### Creating allele-specific ConsHMM annotations

To generate allele specific ConsHMM annotations we used ConsHMM v1.1, containing the new updateInitialParams and ReassignVariantState commands. ConsHMM v1.1 is built on top of ChromHMM v1.20. The updateInitialParams takes as input the parameters of a ConsHMM model and a genome wide segmentation, and outputs an updated parameter set where the initial state parameters are replaced by the genome wide frequency of each state in the segmentation, to better reflect the state assignment prior for a variant at any position in the genome. The ReassignVariantState command takes as input the ConsHMM model outputted by updateInitialParams, a file containing the multiple alignment on which the model is based, and a parameter *W*. The file containing the multiple alignment is in a format processed by the parseMAF command of ConsHMM. The parameter *W* indicates to consider *W* flanking bases upstream and also *W* bases downstream of the allele when computing allele-specific conservation state assignments. We note that since ConsHMM uses an HMM, the state assignment at a position of interest can depend on the observations at neighboring positions.

We investigated the effect of different choices of *W* by first sampling a set of 40,000 common variants from dbSNP(Sherry et al. 2001) that are further than 200kb apart, the segment size previously used with ConsHMM for genome segmentations. We then applied ConsHMM as previously done, but with the alternate allele for those common variants(Arneson and Ernst 2019). We did this with the ConsHMM model for hg38 based on the 100-way vertebrate alignment. We then compared the agreement in the conservation state assignment when we applied ReassignVariantState with values of *W* between 1 and 10 and found that the agreement between the procedures plateaued at 99.6%. The final allele-specific annotations were generated using *W* = 10.

We did this for each possible allele as the base in the center of the window, for every nucleotide in the human genome, and for both the hg19 and hg38 100-way vertebrate alignments. For variants in which the flanking region extends past the beginning or end of chromosomes, the missing bases upstream or downstream of the position of interest were marked as positions where the multiple sequence alignment is empty, which ConsHMM encodes as positions where no species align to the reference species.

### Data and code availability

All alignments we used were obtained from the UCSC genome browser or Ensembl(Herrero et al. 2016; Haeussler et al. 2019). The UCSC multiple sequence alignments listed in **Supplementary Table S1** were downloaded from https://hgdownload.soe.ucsc.edu/downloads.html(Haeussler et al. 2019). The Ensembl multiple sequence alignments listed in **Supplementary Table S1** were downloaded from ftp://ftp.ensembl.org/pub/release-97/maf/ensembl-compara/ and ftp://ftp.ensembl.org/pub/release-75/emf/ensembl-compara/ (Herrero et al. 2016).

SiPhy-omega, SiPhy-pi constrained element calls, and HAR calls were downloaded from https://www.broadinstitute.org/mammals-models/29-mammals-project-supplementary-info(Garber et al. 2009; Lindblad-Toh et al. 2011). Narrowpeak fetal brain DNase I Hypersensitivity Sites with Roadmap identifiers E081 and E082 were downloaded from http://egg2.wustl.edu/roadmap/data/byFileType/peaks/consolidated/narrowPeak/(Road map Epigenomics Consortium et al. 2015). The E081 data was used in **fig. 2** and both E081 and E082 were used in **Supplementary fig. S2**. The mouse pseudogene annotations were obtained from https://www.gencodegenes.org/mouse/.

**Figure 1:**
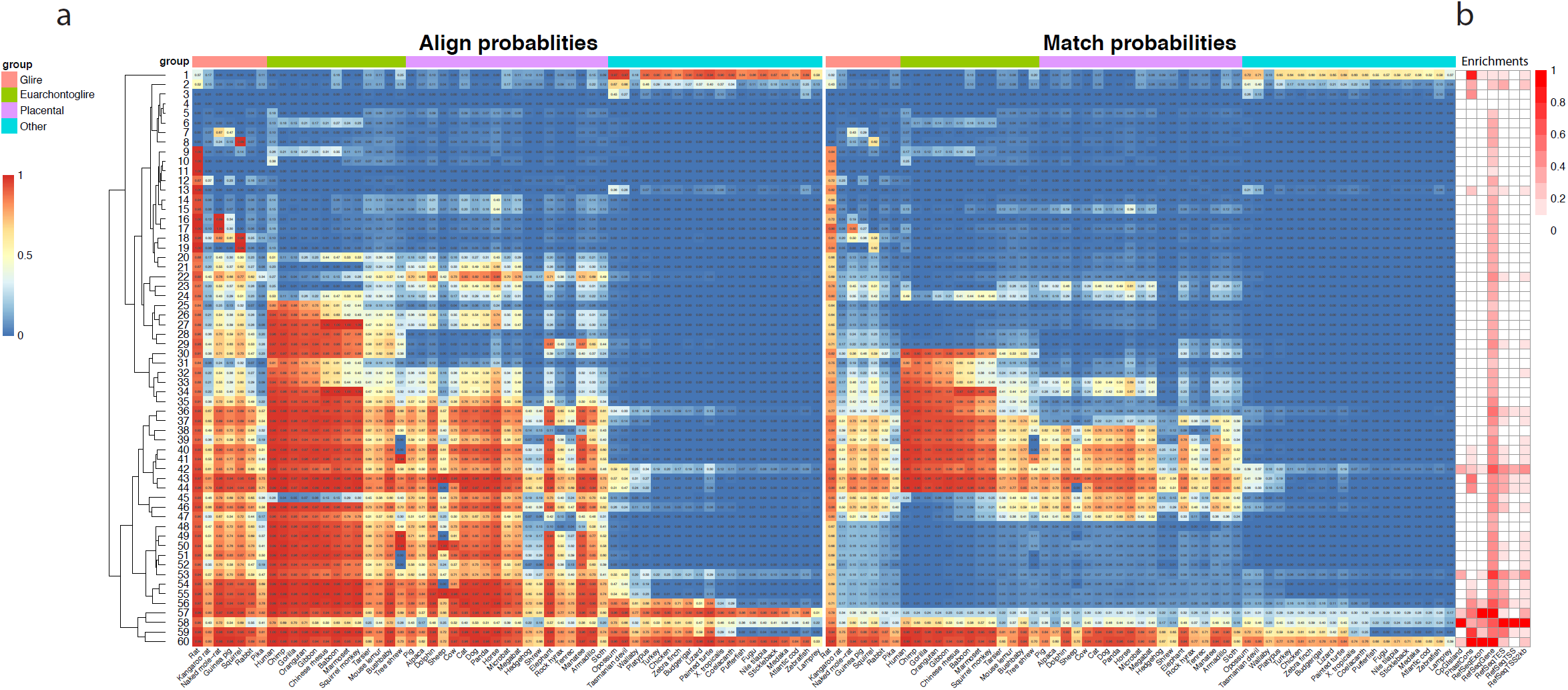
Conservation state emission parameters of a ConsHMM model based on a 60-way alignment of vertebrates to mouse and enrichments for other genomic annotations. **(a)** The rows of the heatmap correspond to conservation states and the columns of the heatmap correspond to species. For each state and species, the left half of the heatmap contains the probability of that species aligning to the mouse genome (mm10) at the position, which means there is a non-indel nucleotide present at the position in the alignment for the species (one minus the probability of the not aligning observation). The right half of the heatmap contains the probability of observing a species matching the mouse genome at the position, which means there is a nucleotide present in the alignment at the position for the species that is the same as in mouse. Species are ordered by phylogenetic distance to mouse and grouped by major clades. States are ordered by by hierarchical clustering, using optimal leaf ordering (Bar-Joseph et al. 2001) implemented in ConsHMM. **(b)** The columns of the heatmap indicate the relative enrichments of conservation states for CpG Islands, PhastCons elements, and RefSeq exons, genes, transcription end sites, transcription start sites, and a 2kb window around transcription start sites (**Materials and Methods**). Each column of the enrichment heatmap was normalized by subtracting the minimum value of the column and dividing by its range. The values in the enrichment heatmap can be found in the **Supplementary Data**.

**Figure 2:**
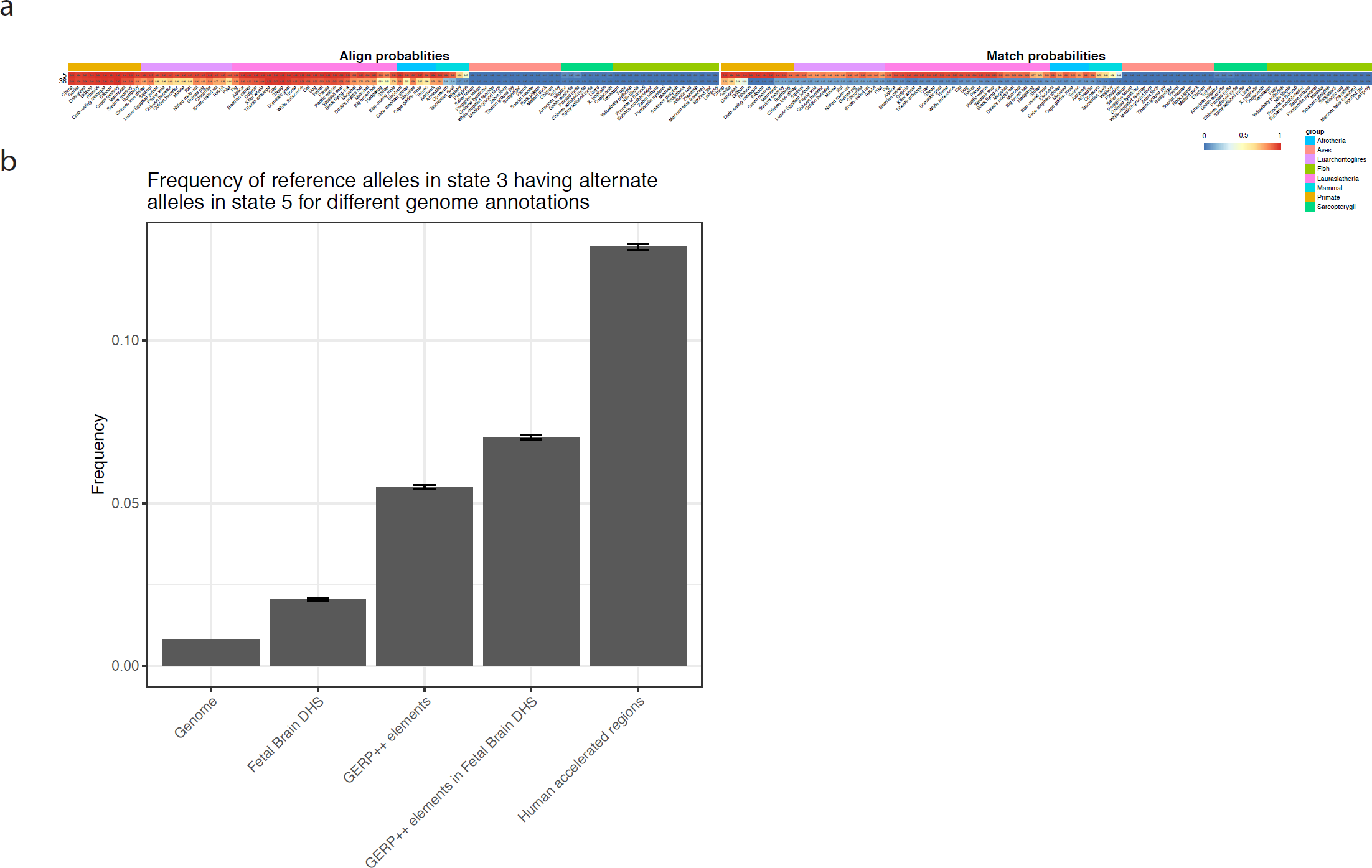
Example of additional information within allele specific conservation state assignments. **(a)** Emission parameters of two states from a 100 state model based on a 100-way vertebrate alignment to the hg38 human genome, states 5 and 36. State 5 is associated with high frequency of aligning and matching through mammals, while state 36 is associated with high frequency of aligning through mammals, but only matching through a subset of primates. The heatmap is structured analogously to the heatmap in **fig. 1**, with the species ordered by phylogenetic distance to human in this case. **(b)** Bar graph showing for different subset of variants assigned to state 36 based on the reference allele, the frequency of assignment change to state 5 out of all possible alternate alleles. The ‘Genome’ category shows this frequency for all variants assigned to state 36 based on the reference allele. The rest of the columns shows the frequency when restricting to subset of those variants positioned in a Fetal Brain DNase I hypersensitivity site (DHS), GERP++ element, the intersection of Fetal Brain DHS and GERP++ elements, and human accelerated regions. Error bars represent a 95% binomial confidence interval computed using a normal approximation of the error around the estimate.

PhastCons constrained element calls, RefSeq and CpG Island annotations and dbSNP v150 variants were obtained from the UCSC genome browser. The ConsHMM model parameters and the corresponding genomic segmentations and annotations are available at http://www.biolchem.ucla.edu/labs/ernst/ConsHMMAtlas. The allele specific state annotations for the human genome and link to the R shiny app can also be found through the same URL. The ConsHMM software is available at https://github.com/ernstlab/ConsHMM.

## Results and Discussion

### ConsHMM annotations for additional organisms and multiple-sequence alignments

We generated an additional 22 ConsHMM genome annotations that in addition to human genome include annotations for the mouse, rat, dog, zebrafish, fruit fly, C. elegans and S. cerevisiae genomes (**Supplementary Table S1, Supplementary Data**). For some species, we generated multiple different genome annotations that corresponded to different sets of species in the multi-species alignment, different alignment methods used to generate the alignment, or different assemblies of the reference genome. We applied ConsHMM as previously described(Arneson and Ernst 2019), except setting the number of states for a model based on the number of species in the alignment (**Materials and Methods**).

We highlight as an illustrative example of one of the new ConsHMM models that we learned, the model for annotating the mouse mm10 genome based on the 60-way Multiz alignment of 59 vertebrates to it (**fig. 1a**). In this model, which has 60 states, ConsHMM identified a number of noteworthy states showing enrichment for other external genomic annotations (**fig. 1b, Supplementary Data**). For example, a state that showed high aligning and matching probabilities in all the species in the alignment, state 60, was the most enriched state for exons (34.9 fold). A different state showed a pattern of moderate probabilities of aligning and matching for almost all species, and showed strong enrichment for CpG islands (50.4 fold) and transcription start sites (34 fold). Another state, state 1, had high aligning probabilities only in distal species to mouse, which is likely capturing alignment artifacts, though still had a 12 fold enrichment for PhastCons constrained element calls. There were three other states (states 2, 3, and 13), which had similar though weaker versions of the state 1 alignment pattern and also enriched for PhastCons elements (3.8-6.4 fold). These four states were all also enriched for pseudogenes, with enrichments ranging from 3.3 to 55.9 fold. These various state patterns and corresponding enrichment were similar to those found for a previously analyzed human conservation state annotation(Arneson and Ernst 2019).

### Allele Specific ConsHMM annotations

Previously, ConsHMM could only generate position specific conservation states based on the allele present in the reference genome. As ConsHMM models the observation of whether the nucleotide present in each other species matches the reference genome, an alternate allele at a position could potentially lead to a very different conservation state assignment. Allele specific annotations could thus be informative to studying genetic variation, but directly applying ConsHMM for every observed variant would not be computationally practical.

To address this challenge we extended ConsHMM to be able to compute conservation state assignments for any alternate allele with high accuracy under two assumptions. The first assumption is that it is sufficient to assume an alternate allele would not cause changes to the multi-species alignment except for the nucleotide present in the reference genome. The second assumption is that it is sufficient to consider a small local window around each variant to derive a state annotation opposed to segmenting 200kb at time as previously done(Arneson and Ernst 2019). We empirically verified this second assumption by considering a range of window sizes upstream and downstream of a variant and showing that a window of size 21 (10 bases upstream and 10 bases downstream) obtained 99.6% agreement in the conservation state assignments compared to applying ConsHMM as previously applied for a set of 40,000 common variants (**Materials and Methods, Supplementary fig. S1**).

Using this extended version of ConsHMM we produce allele specific conservation state annotations for each possible single nucleotide alternate allele for both the hg19 and hg38 human reference genomes based on ConsHMM models trained on 100-way vertebrate alignments. To demonstrate the additional information in having allele specific conservation state assignments beyond just the reference genome we consider the set of positions that were assigned to a state in the human hg38 model associated with high probability of aligning through mammals, but a high probability of matching in a only a few primates, state 36. We then analyzed for different subsets of positions the frequency at which an alternate allele result in a conservation state assignment to a very different state that had high probability of both aligning and matching in many mammals, state 5 (**fig. 2**).

For only 0.8% of possible alternate alleles for reference allele in state 36, we saw the conservation state assignment change to state 5. Interestingly, we saw this percentage increase substantially for subsets of positions with other unique annotations. Among positions in Fetal Brain DNase I hypersensitive sites(Roadmap Epigenomics Consortium et al. 2015), the percentage was 2% and for those in GERP++ constrained elements(Davydov et al. 2010) it was 5% (**fig. 2b**). The percentage increased to 7% for those annotated as both. The percentage increased even further to 12% for previously annotated bases in human accelerated regions (HAR)(Lindblad-Toh et al. 2011). Similar percentages were found when using other sets of conserved elements and another Fetal Brain DNase I hypersensitivity data set (**Supplementary fig. S2**). These results highlight how allele specific conservation state assignments provide additional information beyond the conservation state assignment from the reference allele.

### Web-interface for visualization of parameters and annotation enrichments of ConsHMM models

We created a web interface built on an R shiny app in which one can browse a representation of emission parameters of ConsHMM models and annotation enrichments (**fig. 3**). Users can access the models trained on each of the reference genomes and multiple sequence alignments listed in **Supplementary Table S1**. The app generates an interactive heatmap containing for each state and species, the model probability for that species aligning the reference genome, and also the probability of having a nucleotide matching the human reference. The interface allows a user to select a subset of states and/or species in the alignment to display, for ease of visualization. Lastly, the interface allows users to display precomputed enrichments of states for external annotations. These include enrichments for existing annotations of gene bodies, exons, transcription start and end sites, and the PhastCons elements called on the same alignment, when available.

**Figure 3:**
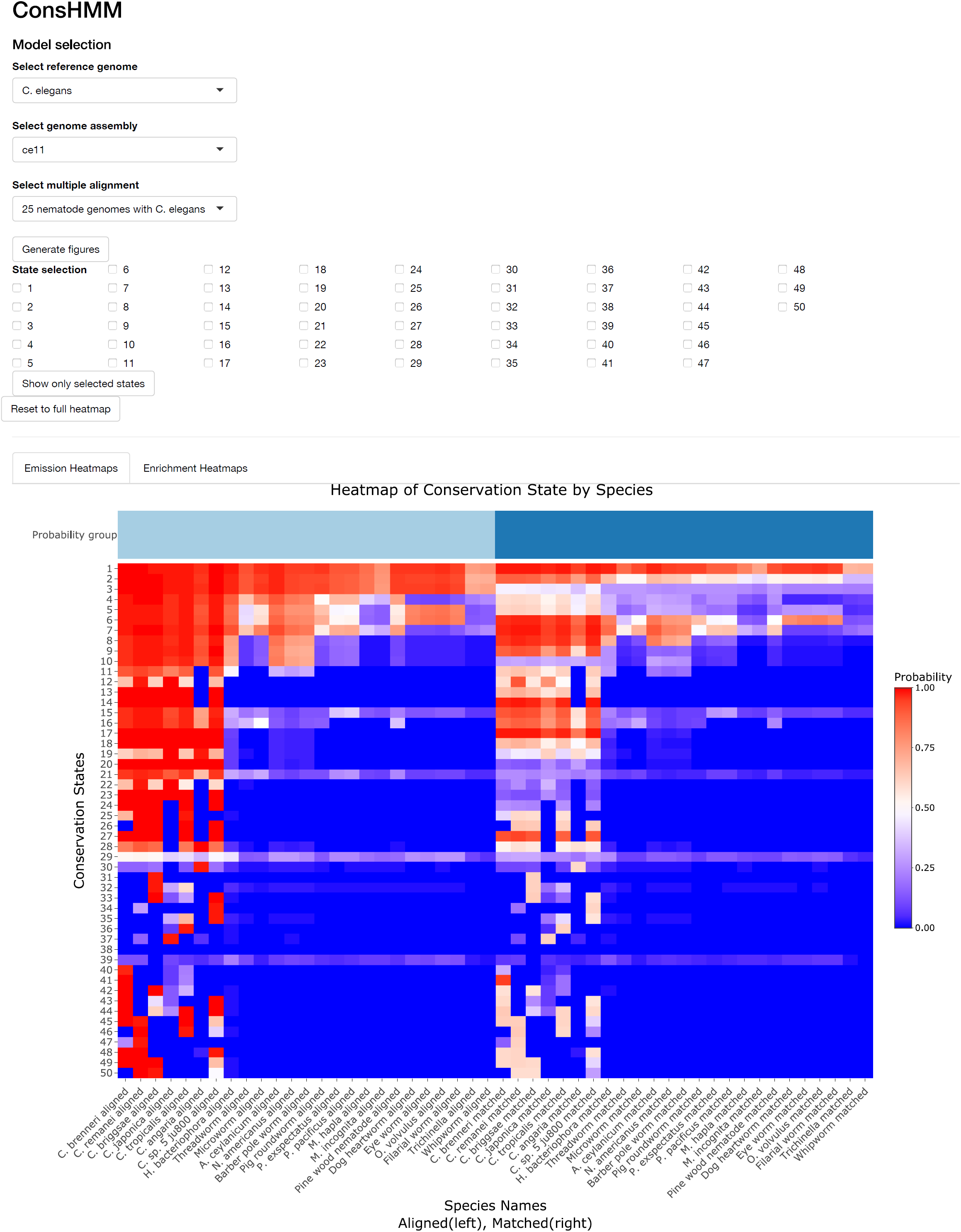
Screenshot of the ConsHMM R Shiny App. The screenshot captures a representation of the emission probabilities of a 50 state model based on a 26-way alignment of nematodes with C. elegans. The dropdown menus at the top of the webpage allows users to select a different reference organism, genome or multiple sequence alignment for which to generate similar heatmaps. Each row in the heatmap corresponds to a state and each column corresponds to a species. As in **fig. 1**., the left half of the heatmap contains the probability of a species aligning the reference genome in the alignment, and the right half of the heatmap contains the probability of a species matching the reference genome in the alignment. The rows are sorted by hierarchical clustering, using optimal leaf ordering (Bar-Joseph et al. 2001) implemented in ConsHMM and the columns are sorted by phylogenetic distance to the reference genome in the alignment. The phylogenetic distance was extracted from the Ensembl Species Tree (Herrero et al. 2016). The checkboxes in the ‘state selection’ area of the app allow users to subset the heatmap to certain states of interest.

## Supporting information

Supplementary emission heatmaps

Supplementary model enrichments

## Acknowledgements

We thank the members of the Ernst lab for useful discussions.

