## Supplementary emission heatmaps for "ConsHMM Atlas: conservation state annotations for major genomes and human genetic variation"

This file contains a representation of ConsHMM parameters as heatmaps for 22 different models.

canFam3 Ensembl 97 EPO 38 mammals

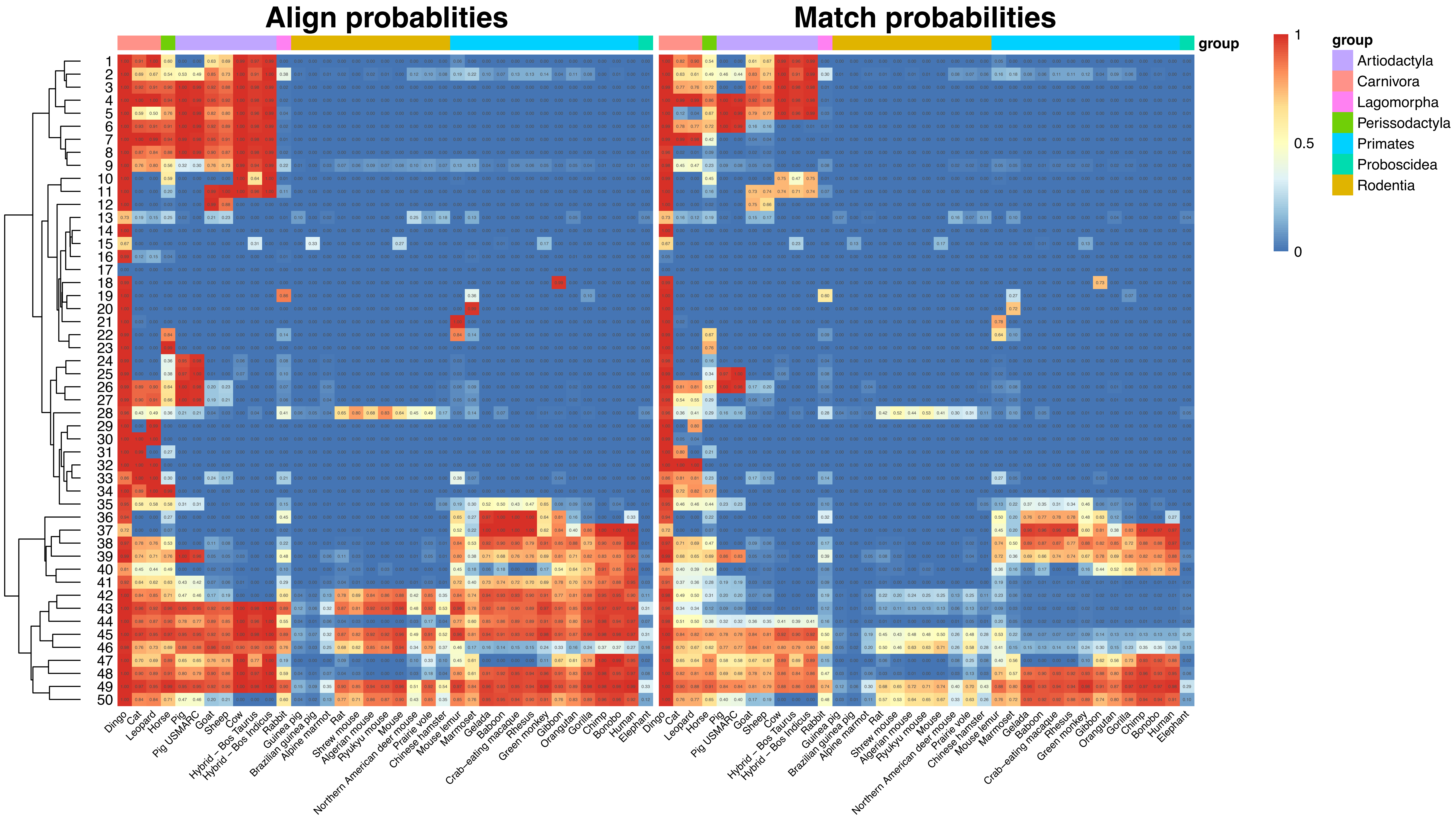

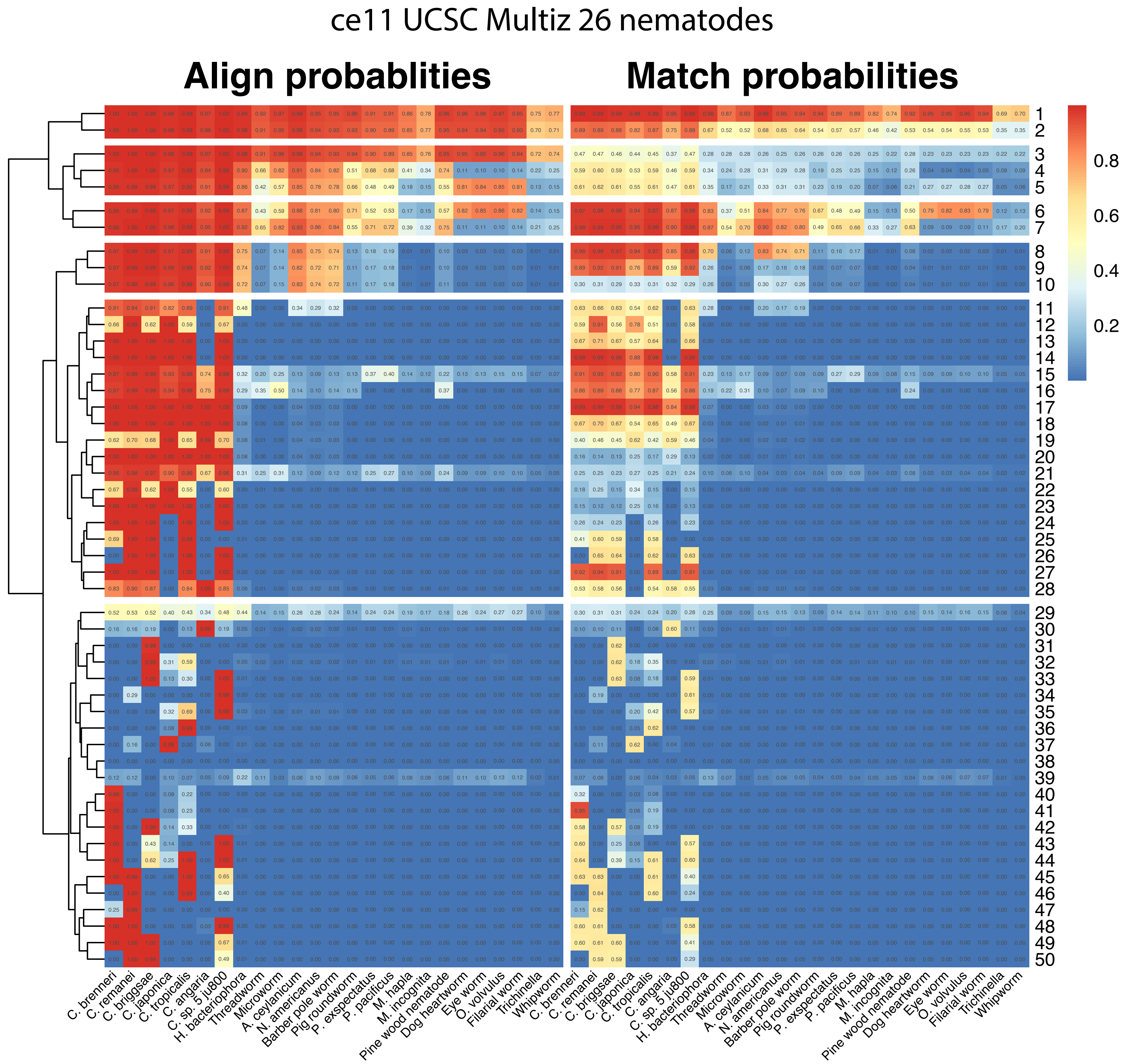

ce11 UCSC Multiz 26 nematodes
Align probablities
Match probabilities
0.8
0.6
0.4
0.2
1 2 3 4 5 6 7 8 9 10 11 12 13 14 15 16 17 18 19 20 21 22 23 24 25 26 27 28 29 30 31 32 33 34 35 36 37 38 39 40 41 42 43 44 45 46 47 48 49 50
C. brenneri
C. remanei
C. briggsae
C. japonica
C. tropicalis
C. angaria
C. sp. 5 ju800
H. bacteriophora
Threadworm
Microworm
A. ceylanicum
N. americanus
Barber pole worm
Pig roundworm
P. exspectatus
P. pacificus
M. hapla
M. incognita
Pine wood nematode
Dog heartworm
Eye worm
O. volvulus
Filarial worm
Trichinella
Whipworm

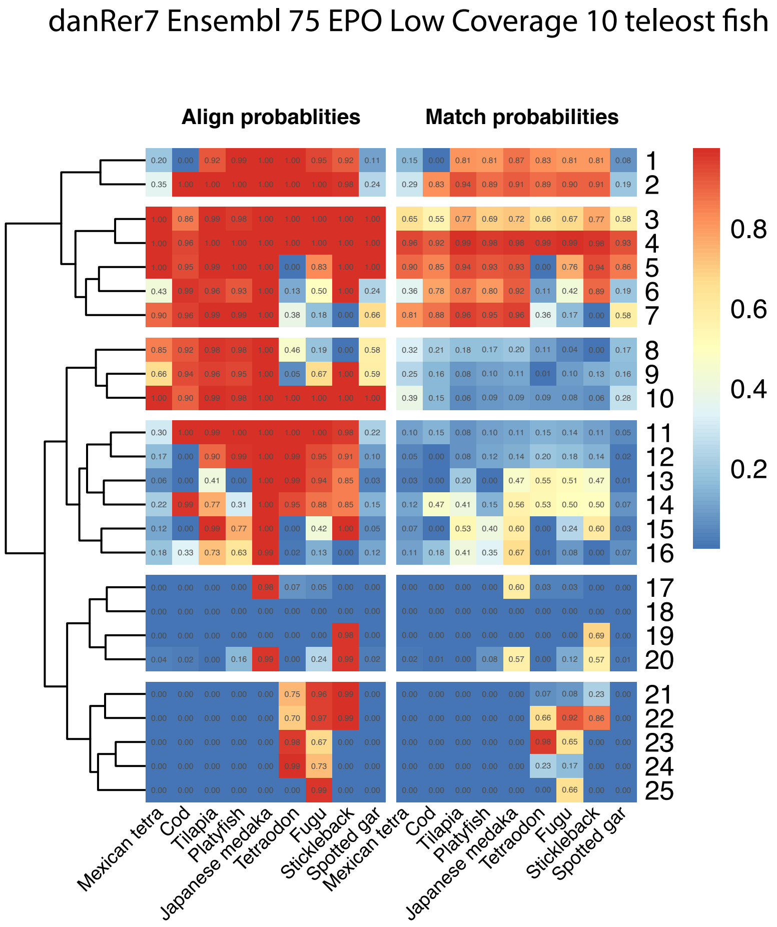
danRer7 Ensembl 75 EPO Low Coverage 10 teleost fish
Align probablities
Match probabilities
1 0.20 0.00 0.92 0.99 1.00 1.00 0.95 0.92 0.11 | 0.15 0.00 0.81 0.81 0.87 0.83 0.81 0.81 0.08
2 0.35 1.00 1.00 1.00 1.00 1.00 1.00 0.98 0.24 | 0.29 0.83 0.94 0.89 0.91 0.89 0.90 0.91 0.19
3 1.00 0.86 0.99 0.98 1.00 1.00 1.00 1.00 1.00 | 0.65 0.55 0.77 0.69 0.72 0.66 0.67 0.77 0.58
4 1.00 0.96 1.00 1.00 1.00 1.00 1.00 1.00 1.00 | 0.96 0.92 0.99 0.98 0.98 0.99 0.99 0.98 0.93
5 1.00 0.95 0.99 1.00 1.00 0.00 0.83 1.00 1.00 | 0.90 0.85 0.94 0.93 0.93 0.00 0.76 0.94 0.86
6 0.43 0.99 0.96 0.93 1.00 0.13 0.50 1.00 0.24 | 0.36 0.78 0.87 0.80 0.92 0.11 0.42 0.89 0.19
7 0.90 0.96 0.99 0.99 1.00 0.38 0.18 0.00 0.66 | 0.81 0.88 0.96 0.95 0.96 0.36 0.17 0.00 0.58
8 0.85 0.92 0.98 0.98 1.00 0.46 0.19 0.00 0.58 | 0.32 0.21 0.18 0.17 0.20 0.11 0.04 0.00 0.17
9 0.66 0.94 0.96 0.95 1.00 0.05 0.67 1.00 0.59 | 0.25 0.16 0.08 0.10 0.11 0.01 0.10 0.13 0.16
10 1.00 0.90 0.99 0.98 1.00 1.00 1.00 1.00 1.00 | 0.39 0.15 0.06 0.09 0.09 0.09 0.08 0.06 0.28
11 0.30 1.00 0.99 1.00 1.00 1.00 1.00 0.98 0.22 | 0.10 0.15 0.08 0.10 0.11 0.15 0.14 0.11 0.05
12 0.17 0.00 0.90 0.99 1.00 0.99 0.95 0.91 0.10 | 0.05 0.00 0.08 0.12 0.14 0.20 0.18 0.14 0.02
13 0.06 0.00 0.41 0.00 1.00 0.99 0.94 0.85 0.03 | 0.03 0.00 0.20 0.00 0.47 0.55 0.51 0.47 0.01
14 0.22 0.99 0.77 0.31 1.00 0.95 0.88 0.85 0.15 | 0.12 0.47 0.41 0.15 0.56 0.53 0.50 0.50 0.07
15 0.12 0.00 0.99 0.77 1.00 0.00 0.42 1.00 0.05 | 0.07 0.00 0.53 0.40 0.60 0.00 0.24 0.60 0.03
16 0.18 0.33 0.73 0.63 0.99 0.02 0.13 0.00 0.12 | 0.11 0.18 0.41 0.35 0.67 0.01 0.08 0.00 0.07
17 0.00 0.00 0.00 0.00 0.98 0.07 0.05 0.00 0.00 | 0.00 0.00 0.00 0.00 0.60 0.03 0.03 0.00 0.00
18 0.00 0.00 0.00 0.00 0.00 0.00 0.00 0.00 0.00 | 0.00 0.00 0.00 0.00 0.00 0.00 0.00 0.00 0.00
19 0.00 0.00 0.00 0.00 0.00 0.00 0.00 0.98 0.00 | 0.00 0.00 0.00 0.00 0.00 0.00 0.00 0.69 0.00
20 0.04 0.02 0.00 0.16 0.99 0.00 0.24 0.99 0.02 | 0.02 0.01 0.00 0.08 0.57 0.00 0.12 0.57 0.01
21 0.00 0.00 0.00 0.00 0.00 0.75 0.96 0.99 0.00 | 0.00 0.00 0.00 0.00 0.00 0.07 0.08 0.23 0.00
22 0.00 0.00 0.00 0.00 0.00 0.70 0.97 0.99 0.00 | 0.00 0.00 0.00 0.00 0.00 0.66 0.92 0.86 0.00
23 0.00 0.00 0.00 0.00 0.00 0.98 0.67 0.00 0.00 | 0.00 0.00 0.00 0.00 0.00 0.98 0.65 0.00 0.00
24 0.00 0.00 0.00 0.00 0.00 0.99 0.73 0.00 0.00 | 0.00 0.00 0.00 0.00 0.00 0.23 0.17 0.00 0.00
25 0.00 0.00 0.00 0.00 0.00 0.00 0.99 0.00 0.00 | 0.00 0.00 0.00 0.00 0.00 0.00 0.66 0.00 0.00
Mexican tetra
Cod
Tilapia
Platyfish
Japanese medaka
Tetraodon
Fugu
Stickleback
Spotted gar
Mexican tetra
Cod
Tilapia
Platyfish
Japanese medaka
Tetraodon
Fugu
Stickleback
Spotted gar
0.8
0.6
0.4
0.2

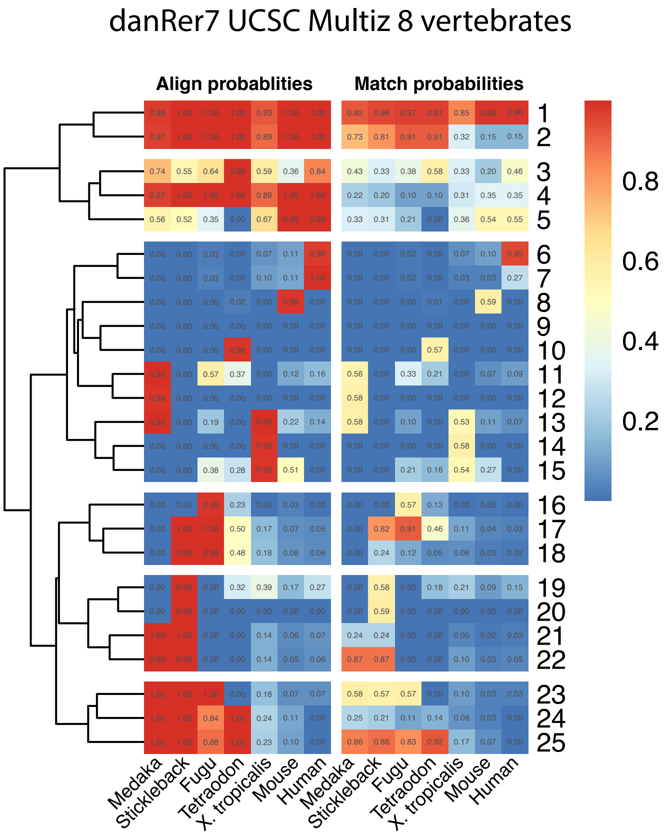
danRer7 UCSC Multiz 8 vertebrates
Align probablities
Match probabilities
0.8
0.6
0.4
0.2
Medaka
Stickleback
Fugu
Tetraodon
X. tropicalis
Mouse
Human
Medaka
Stickleback
Fugu
Tetraodon
X. tropicalis
Mouse
Human

danRer11 Ensembl 97 EPO 25 fish

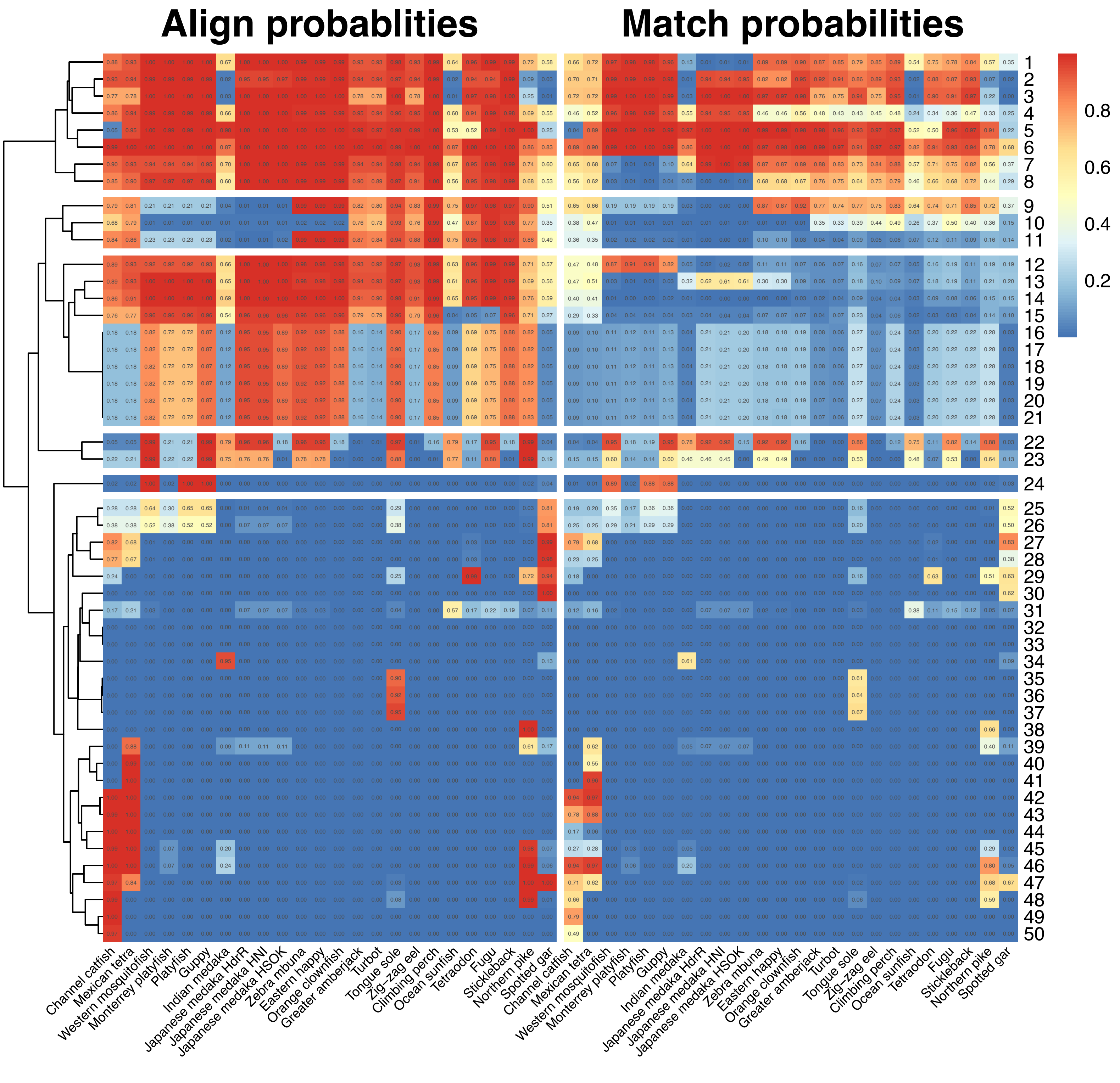

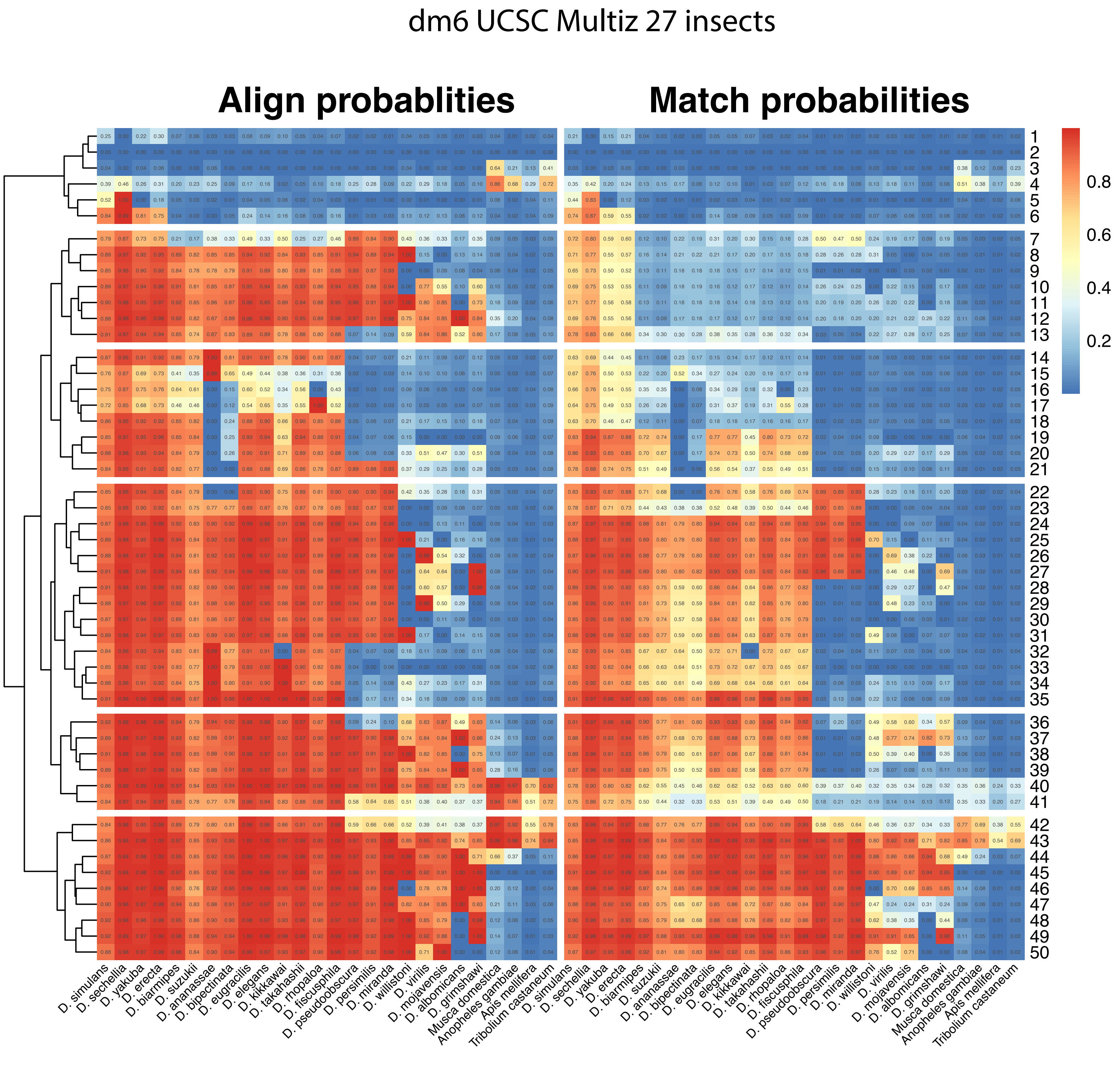

dm6 UCSC Multiz 27 insects
Align probablities
Match probabilities
0.8
0.6
0.4
0.2
D. simulans
D. sechellia
D. yakuba
D. erecta
D. biarmipes
D. suzukii
D. ananassae
D. bipectinata
D. eugracilis
D. elegans
D. kikkawai
D. takahashii
D. rhopaloa
D. ficusphila
D. pseudoobscura
D. persimilis
D. miranda
D. willistoni
D. virilis
D. mojavensis
D. albomicans
D. grimshawi
Musca domestica
Anopheles gambiae
Apis mellifera
Tribolium castaneum

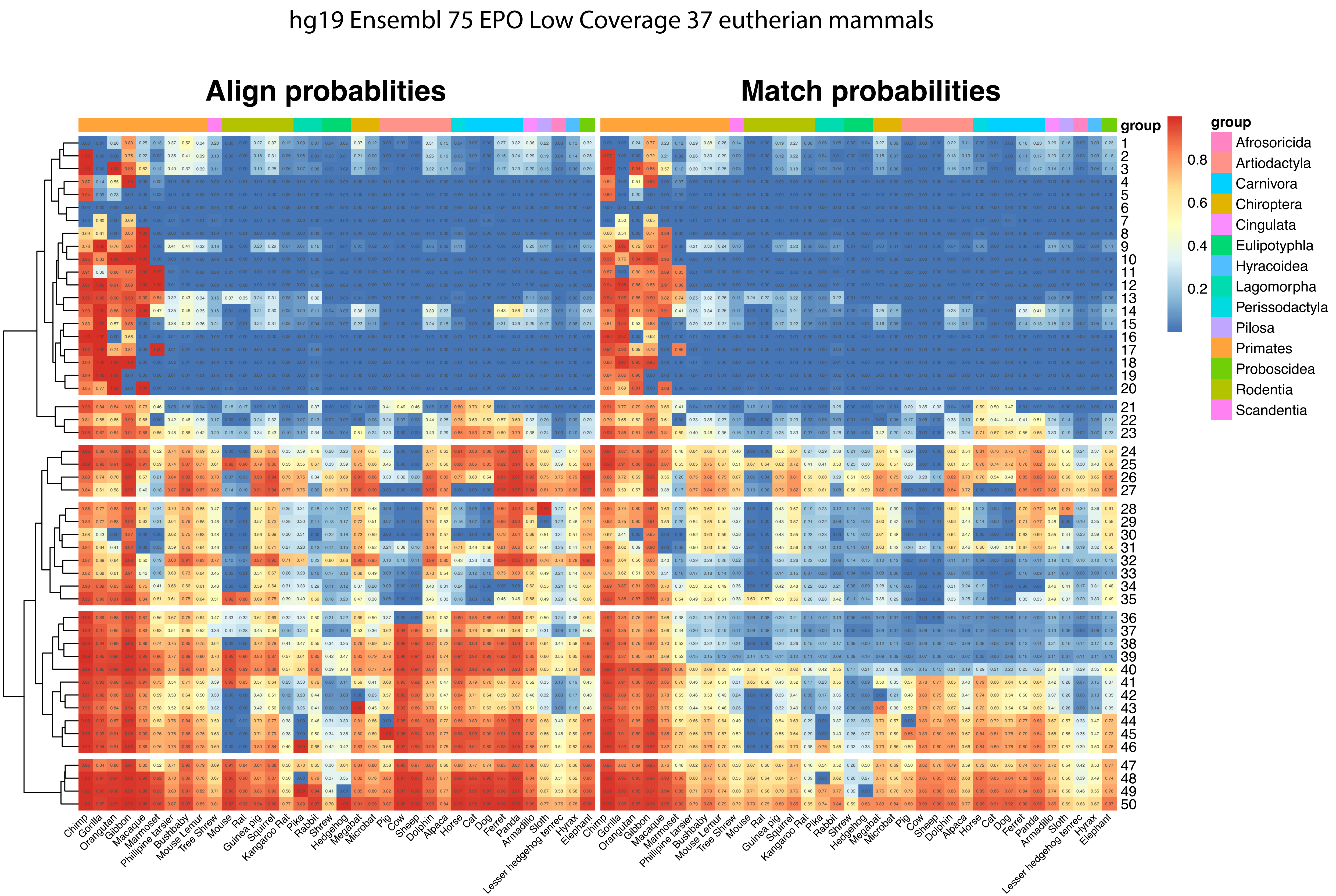
hg19 Ensembl 75 EPO Low Coverage 37 eutherian mammals
Align probablities
Match probabilities
group
Afrosoricida
Artiodactyla
Carnivora
Chiroptera
Cingulata
Eulipotyphla
Hyracoidea
Lagomorpha
Perissodactyla
Pilosa
Primates
Proboscidea
Rodentia
Scandentia
0.8
0.6
0.4
0.2
Chimp
Gorilla
Orangutan
Gibbon
Macaque
Marmoset
Phillipine tarsier
Bushbaby
Mouse Lemur
Tree Shrew
Mouse
Rat
Guinea pig
Squirrel
Kangaroo Rat
Pika
Rabbit
Shrew
Hedgehog
Megabat
Microbat
Pig
Cow
Sheep
Dolphin
Alpaca
Horse
Cat
Dog
Ferret
Panda
Armadillo
Sloth
Lesser hedgehog tenrec
Hyrax
Elephant

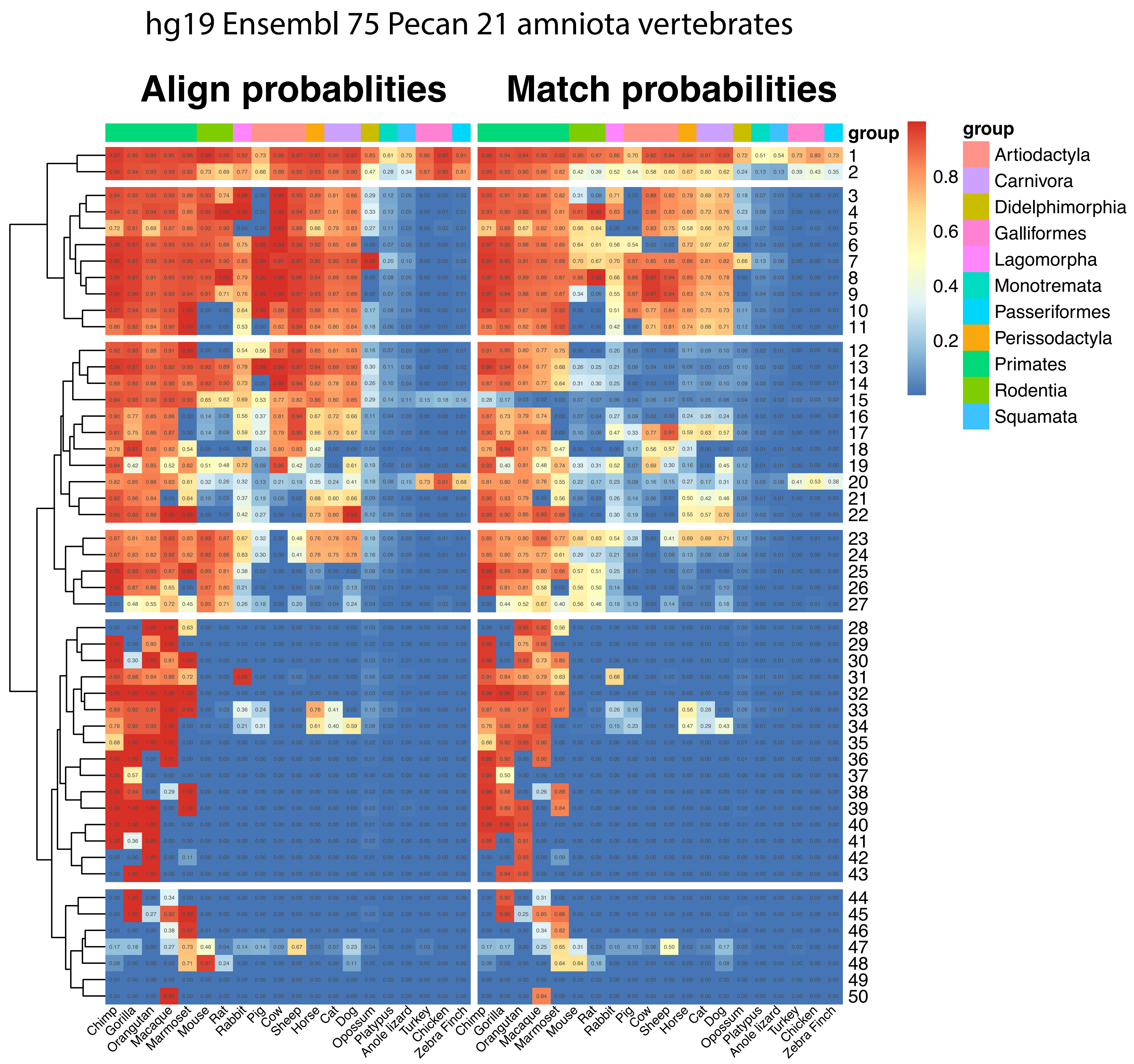
hg19 Ensembl 75 Pecan 21 amniota vertebrates
Align probablities
Match probabilities
group
1 2 3 4 5 6 7 8 9 10 11 12 13 14 15 16 17 18 19 20 21 22 23 24 25 26 27 28 29 30 31 32 33 34 35 36 37 38 39 40 41 42 43 44 45 46 47 48 49 50
Chimp Gorilla Orangutan Macaque Marmoset Mouse Rat Rabbit Pig Cow Sheep Horse Cat Dog Opossum Platypus Anole lizard Turkey Chicken Zebra Finch
0.8
0.6
0.4
0.2
group
Artiodactyla
Carnivora
Didelphimorphia
Galliformes
Lagomorpha
Monotremata
Passeriformes
Perissodactyla
Primates
Rodentia
Squamata

hg38 Ensembl 97 EPO Low Coverage 91 eutherian mammals

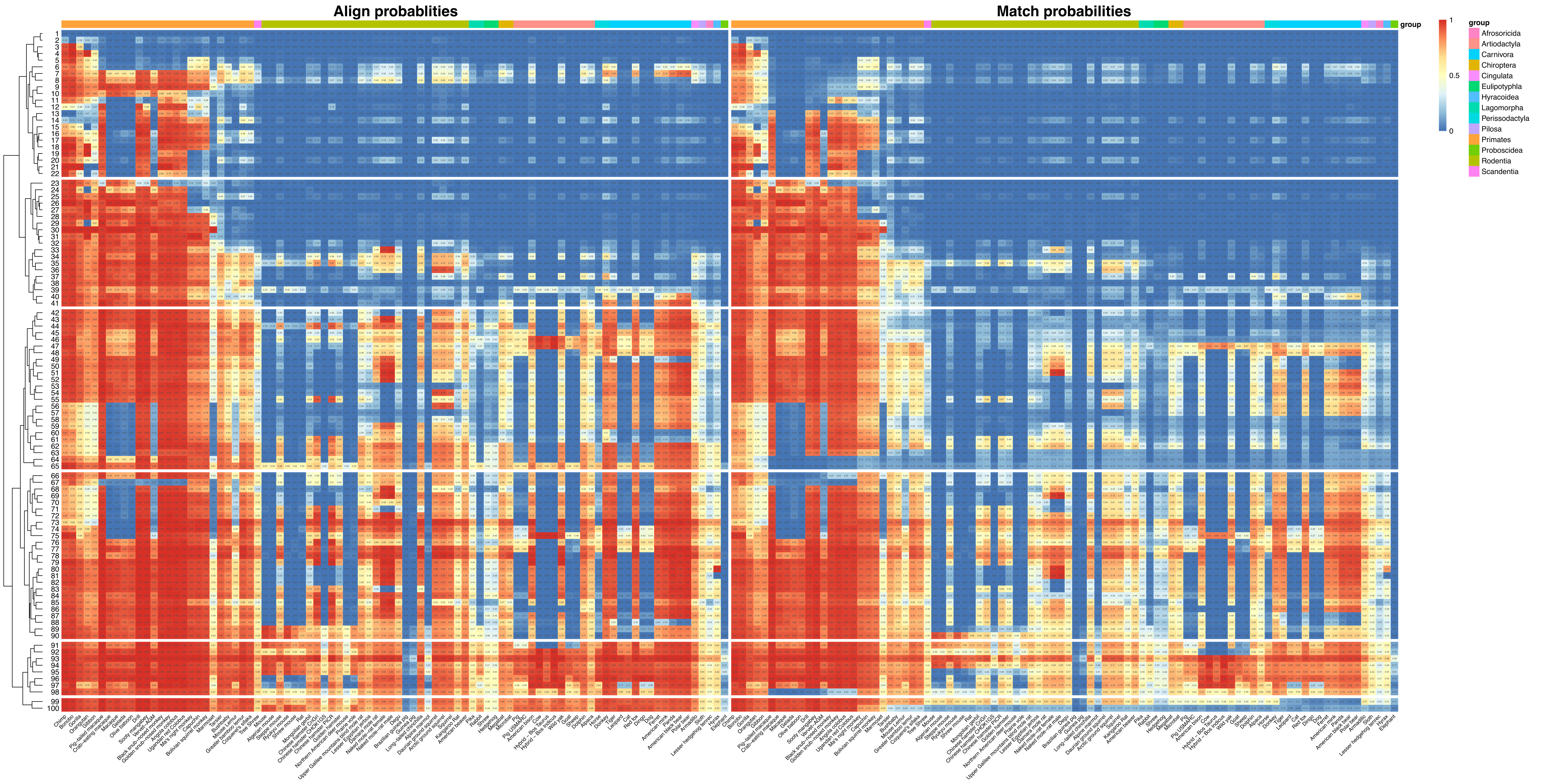

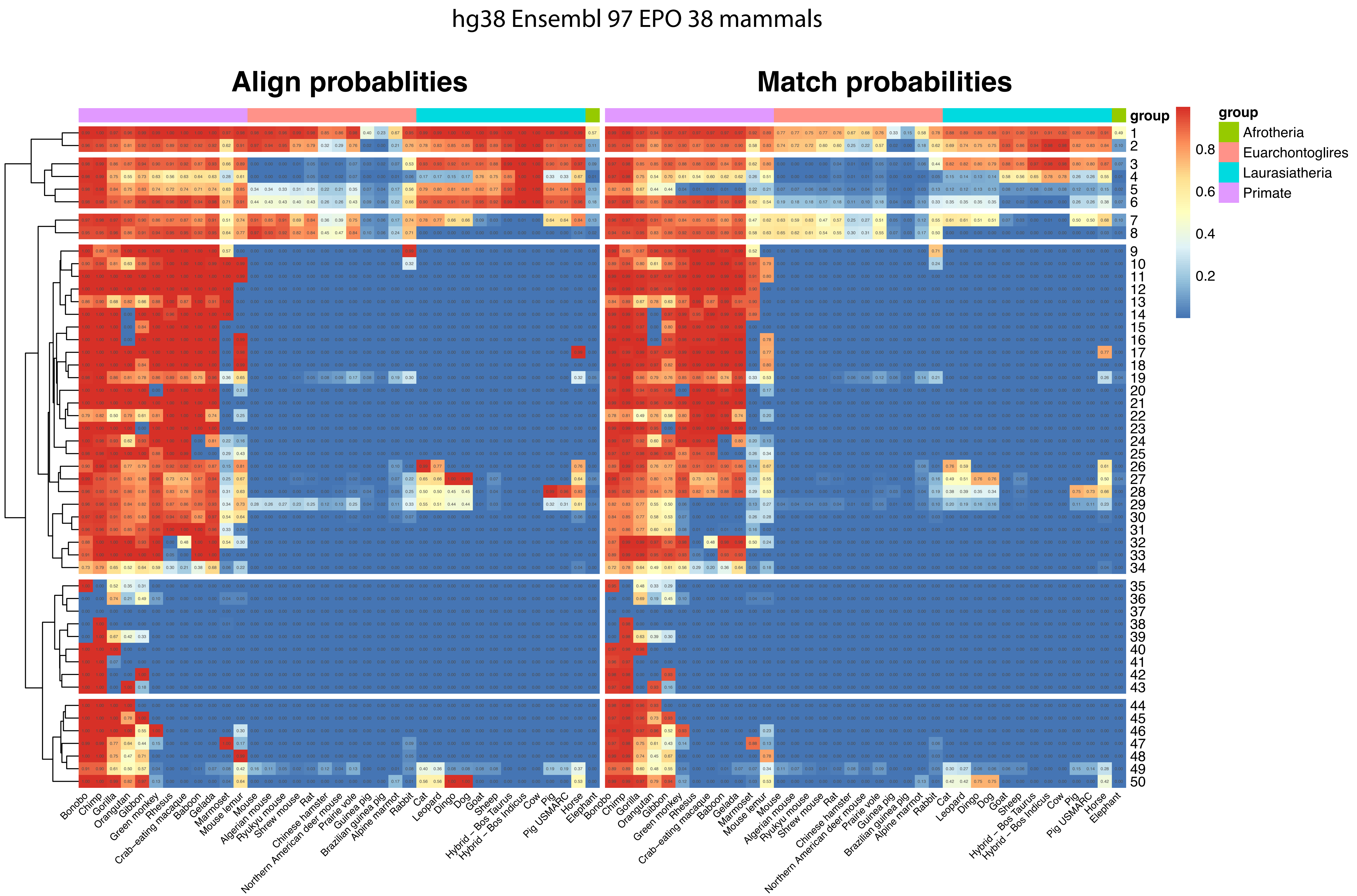
hg38 Ensembl 97 EPO 38 mammals
Align probablities
Match probabilities
group
Afrotheria
Euarchontoglires
Laurasiatheria
Primate
0.8
0.6
0.4
0.2
Bonobo
Chimp
Gorilla
Orangutan
Gibbon
Green monkey
Rhesus
Crab−eating macaque
Baboon
Gelada
Marmoset
Mouse lemur
Mouse
Algerian mouse
Ryukyu mouse
Shrew mouse
Rat
Chinese hamster
Northern American deer mouse
Prairie vole
Guinea pig
Brazilian guinea pig
Alpine marmot
Rabbit
Cat
Leopard
Dingo
Dog
Goat
Sheep
Hybrid − Bos Taurus
Hybrid − Bos Indicus
Cow
Pig
Pig USMARC
Horse
Elephant

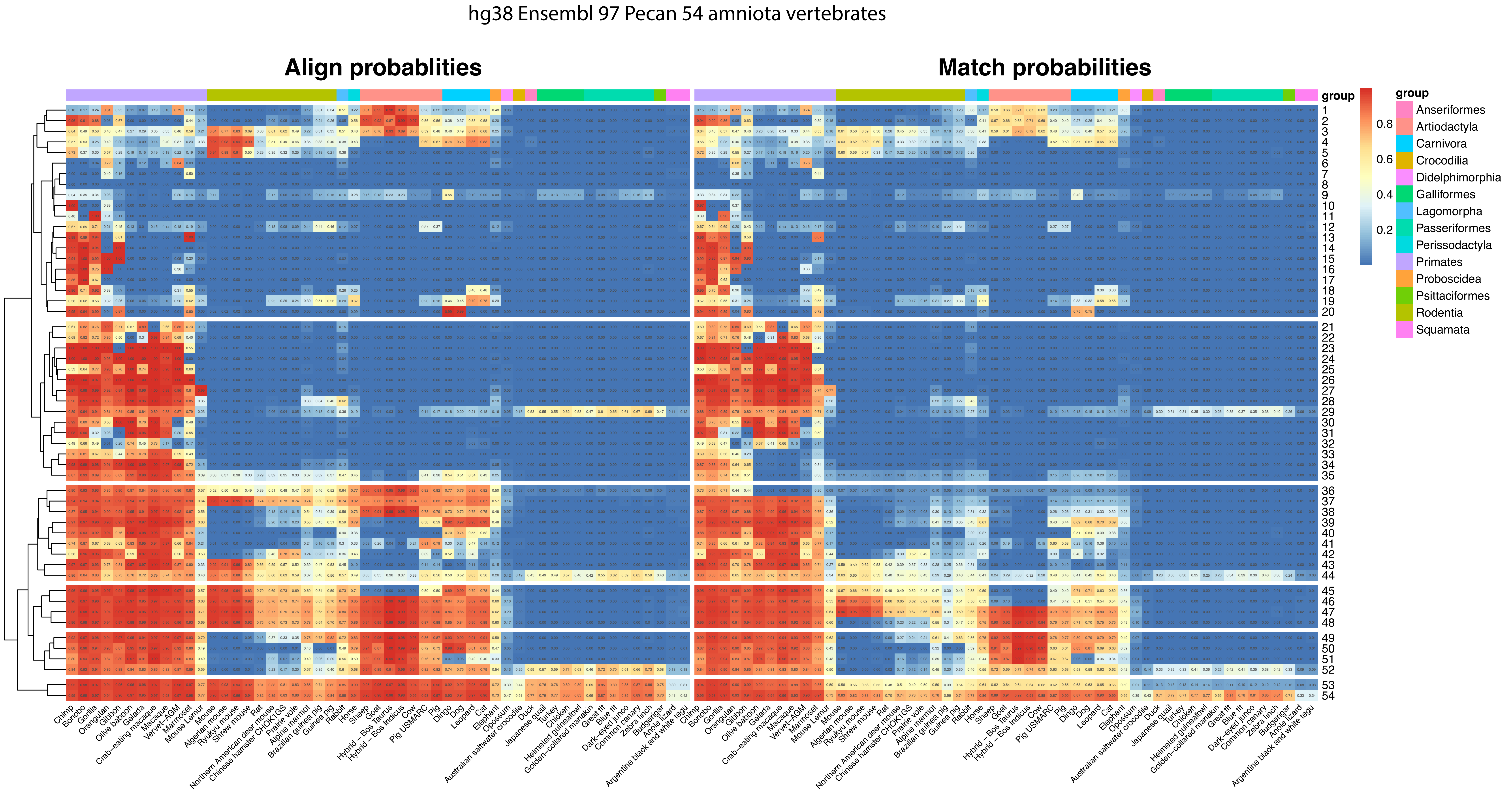
hg38 Ensembl 97 Pecan 54 amniota vertebrates
Align probablities
Match probabilities
group
0.8
0.6
0.4
0.2
group
Anseriformes
Artiodactyla
Carnivora
Crocodilia
Didelphimorphia
Galliformes
Lagomorpha
Passeriformes
Perissodactyla
Primates
Proboscidea
Psittaciformes
Rodentia
Squamata
Chimp
Bonobo
Gorilla
Orangutan
Gibbon
Olive baboon
Gelada
Crab-eating macaque
Macaque
Vervet-AGM
Marmoset
Mouse Lemur
Mouse
Algerian mouse
Ryukyu mouse
Shrew mouse
Rat
Northern American deer mouse
Chinese hamster CHOK1GS
Prairie vole
Alpine marmot
Brazilian guinea pig
Guinea pig
Rabbit
Horse
Sheep
Goat
Hybrid - Bos Taurus
Hybrid - Bos Indicus
Cow
Pig USMARC
Pig
Dingo
Dog
Leopard
Cat
Elephant
Opossum
Australian saltwater crocodile
Duck
Japanese quail
Turkey
Chicken
Helmeted guineafowl
Golden-collared manakin
Great tit
Blue tit
Dark-eyed junco
Common canary
Zebra finch
Budgerigar
Anole lizard
Argentine black and white tegu

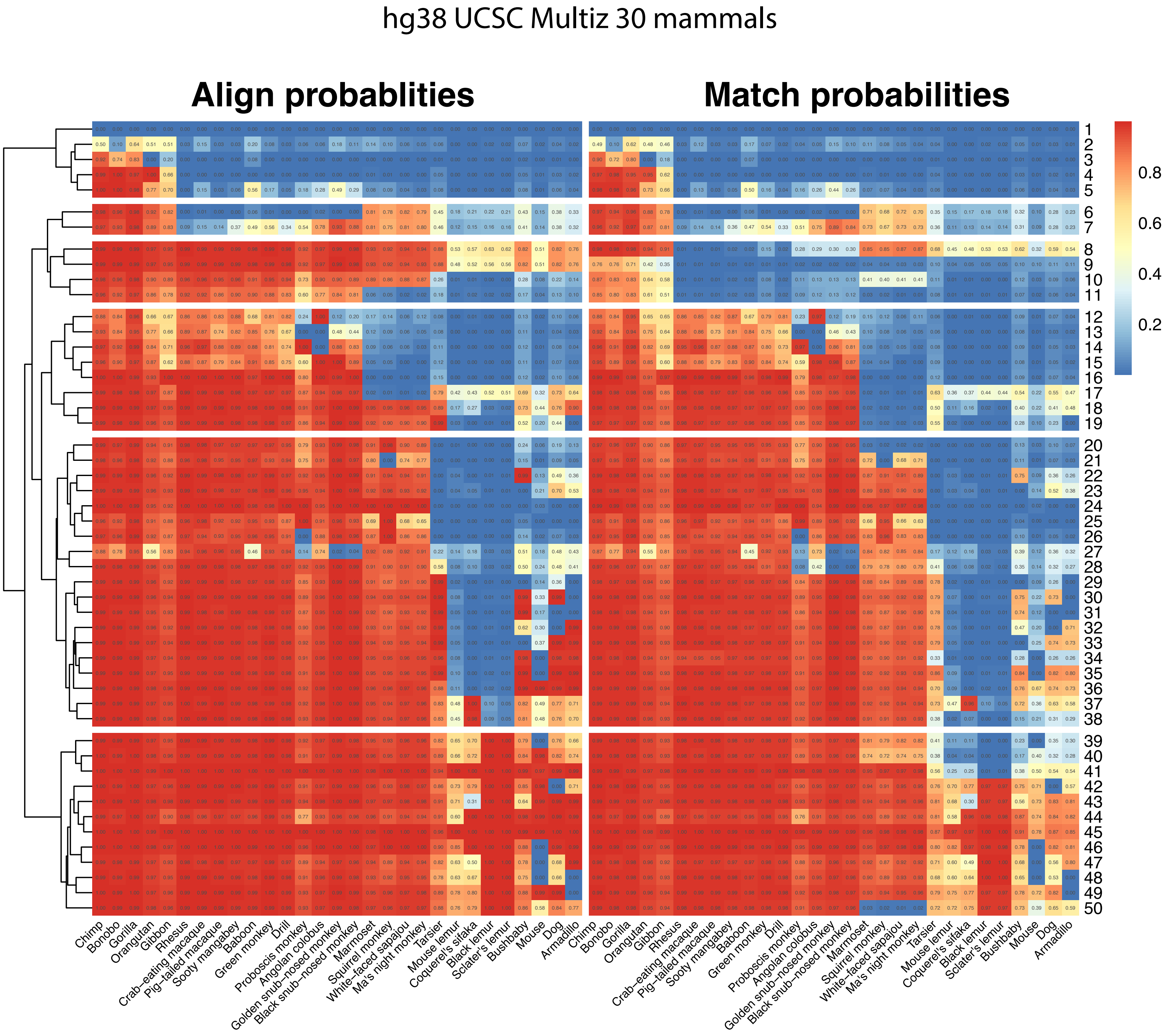
hg38 UCSC Multiz 30 mammals
Align probablities
Match probabilities
0.8
0.6
0.4
0.2
Chimp
Bonobo
Gorilla
Orangutan
Gibbon
Rhesus
Crab-eating macaque
Pig-tailed macaque
Sooty mangabey
Baboon
Green monkey
Drill
Proboscis monkey
Angolan colobus
Golden snub-nosed monkey
Black snub-nosed monkey
Marmoset
Squirrel monkey
White-faced sapajou
Ma's night monkey
Tarsier
Mouse lemur
Coquerel's sifaka
Black lemur
Sclater's lemur
Bushbaby
Mouse
Dog
Armadillo

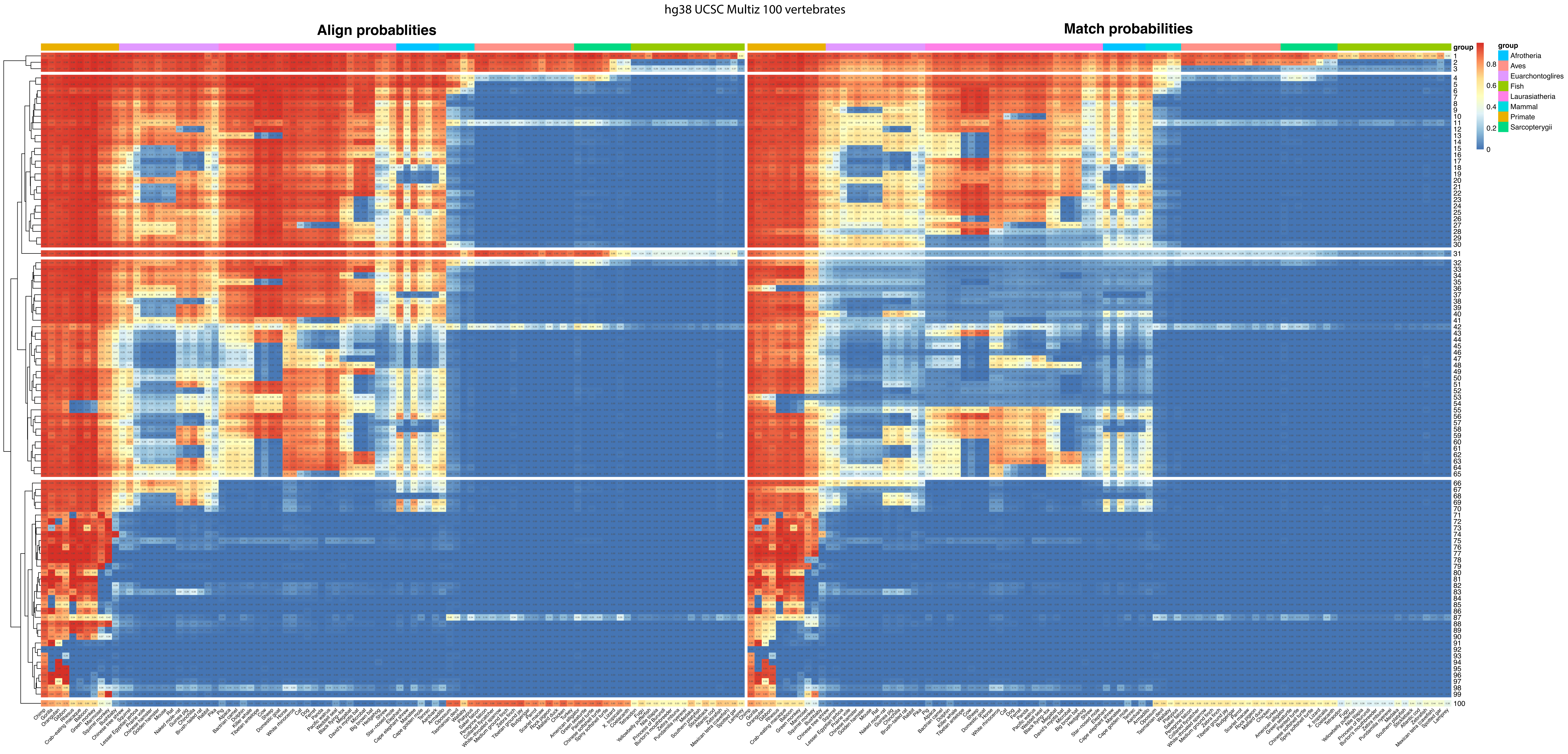
hg38 UCSC Multiz 100 vertebrates
Align probablities
Match probabilities
group
Afrotheria
Aves
Euarchontoglires
Fish
Laurasiatheria
Mammal
Primate
Sarcopterygii
0.8
0.6
0.4
0.2
0

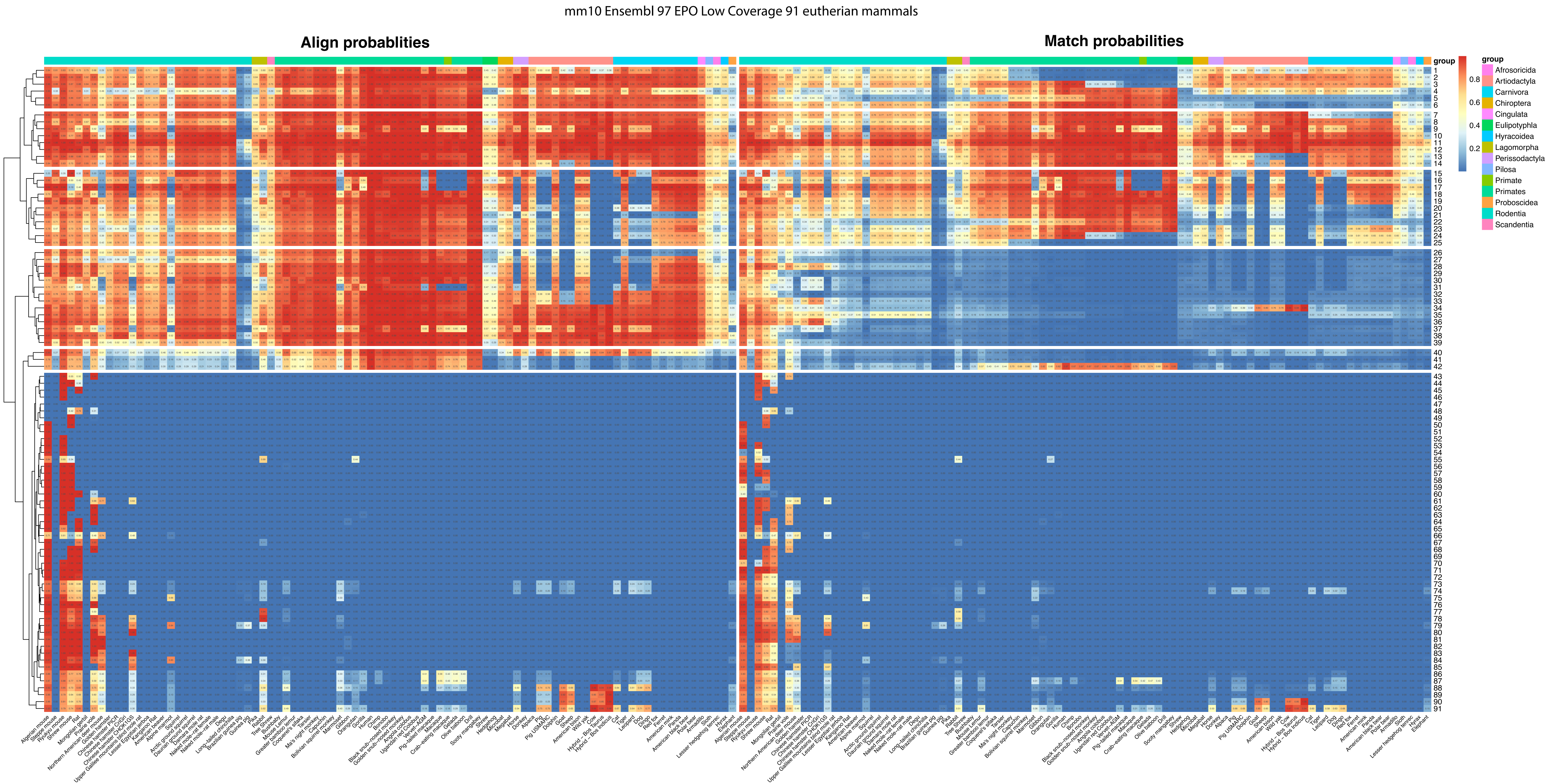
mm10 Ensembl 97 EPO Low Coverage 91 eutherian mammals
Align probablities
Match probabilities
group
Afrosoricida
Artiodactyla
Carnivora
Chiroptera
Cingulata
Eulipotyphla
Hyracoidea
Lagomorpha
Perissodactyla
Pilosa
Primate
Primates
Proboscidea
Rodentia
Scandentia

mm10 Ensembl 97 EPO 38 mammals

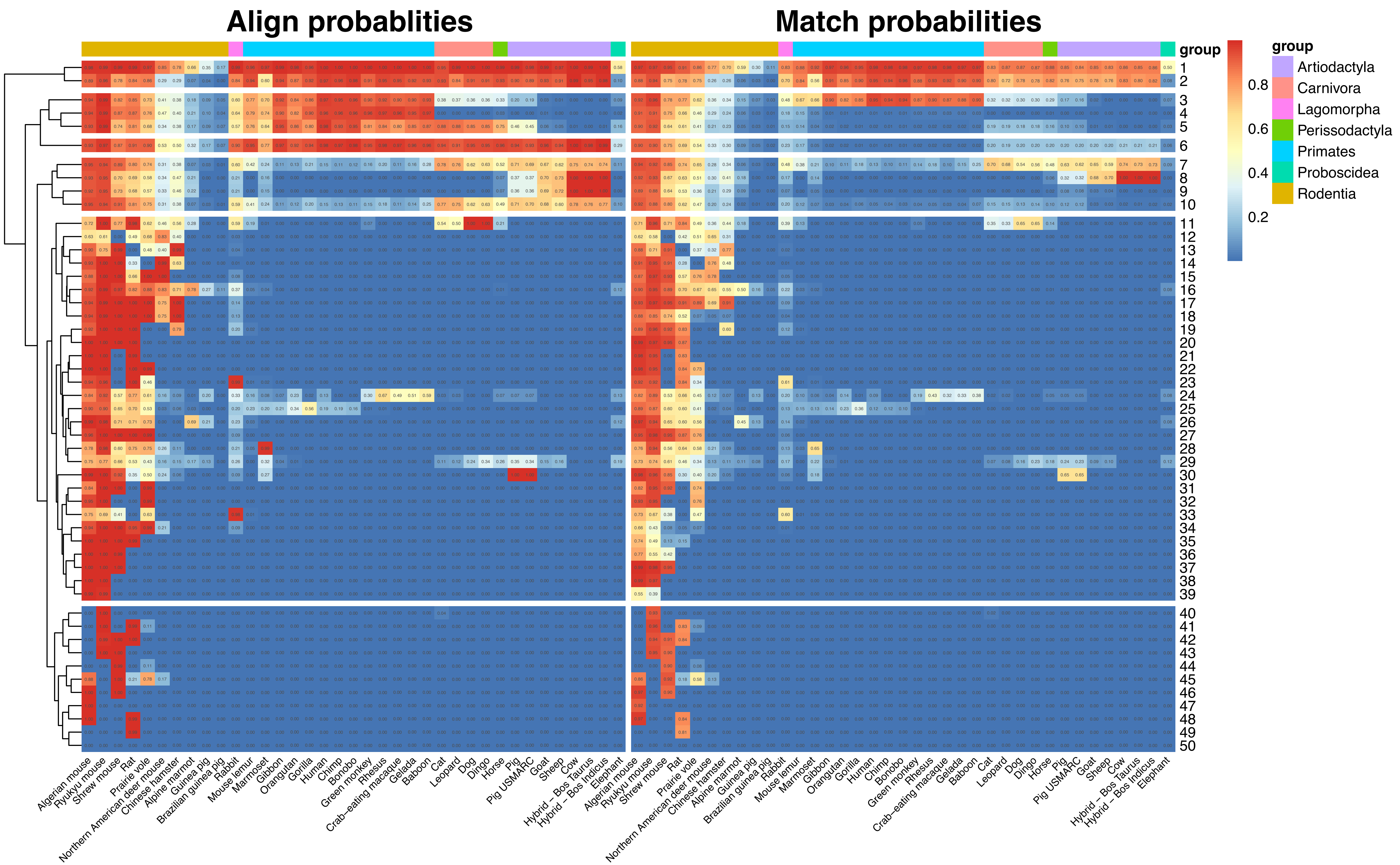

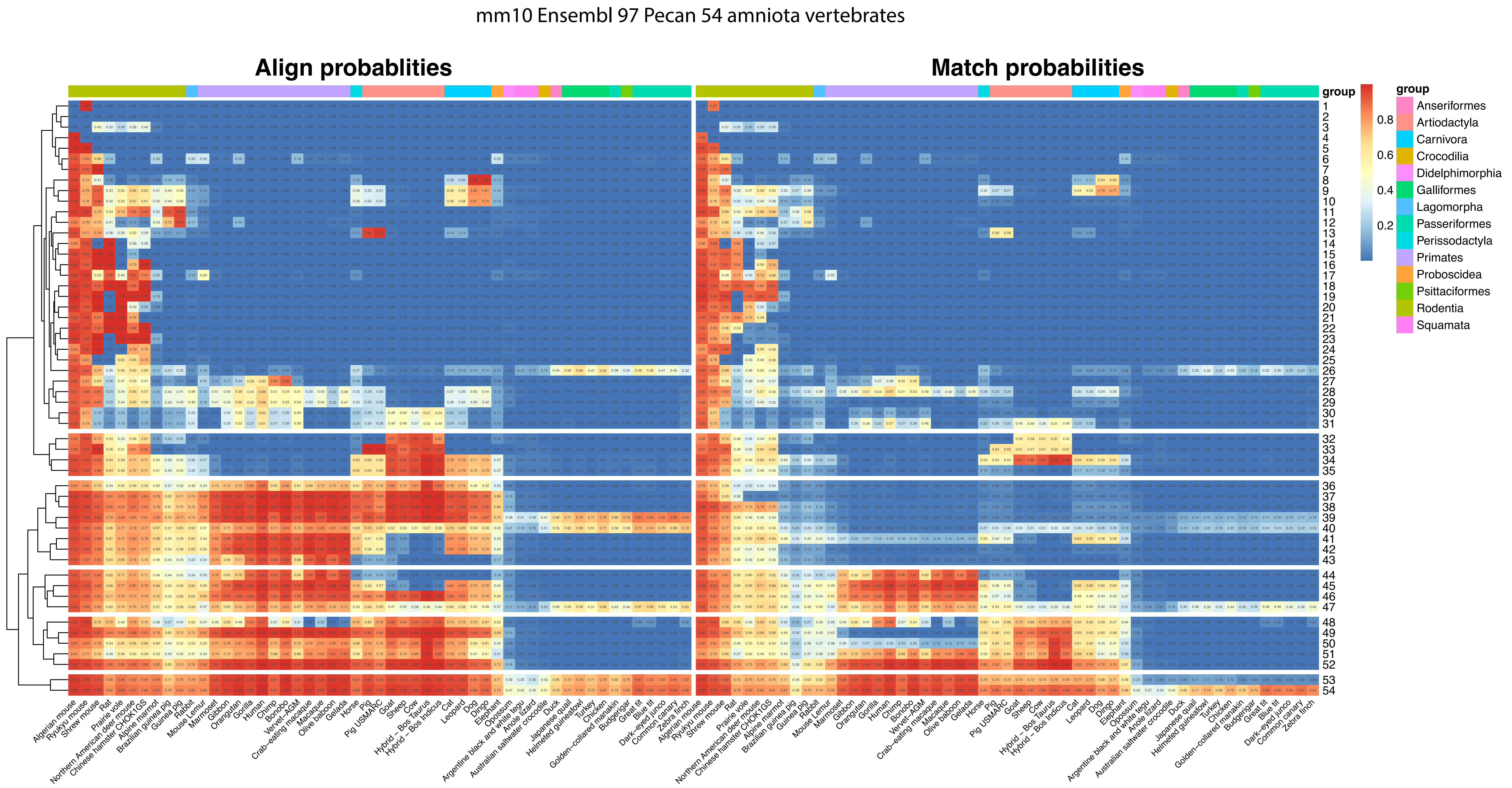
mm10 Ensembl 97 Pecan 54 amniota vertebrates
Align probablities
Match probabilities
group
Anseriformes
Artiodactyla
Carnivora
Crocodilia
Didelphimorphia
Galliformes
Lagomorpha
Passeriformes
Perissodactyla
Primates
Proboscidea
Psittaciformes
Rodentia
Squamata
0.8
0.6
0.4
0.2
Algerian mouse
Ryukyu mouse
Shrew mouse
Rat
Prairie vole
Northern American deer mouse
Chinese hamster CHOK1GS
Alpine marmot
Brazilian guinea pig
Guinea pig
Rabbit
Mouse Lemur
Marmoset
Gibbon
Orangutan
Gorilla
Human
Chimp
Bonobo
Vervet−AGM
Crab−eating macaque
Macaque
Olive baboon
Gelada
Horse
Pig
Pig USMARC
Goat
Sheep
Cow
Hybrid − Bos Taurus
Hybrid − Bos Indicus
Cat
Leopard
Dog
Dingo
Elephant
Opossum
Argentine black and white tegu
Anole lizard
Australian saltwater crocodile
Duck
Japanese quail
Helmeted guineafowl
Turkey
Chicken
Golden−collared manakin
Budgerigar
Great tit
Blue tit
Dark−eyed junco
Common canary
Zebra finch

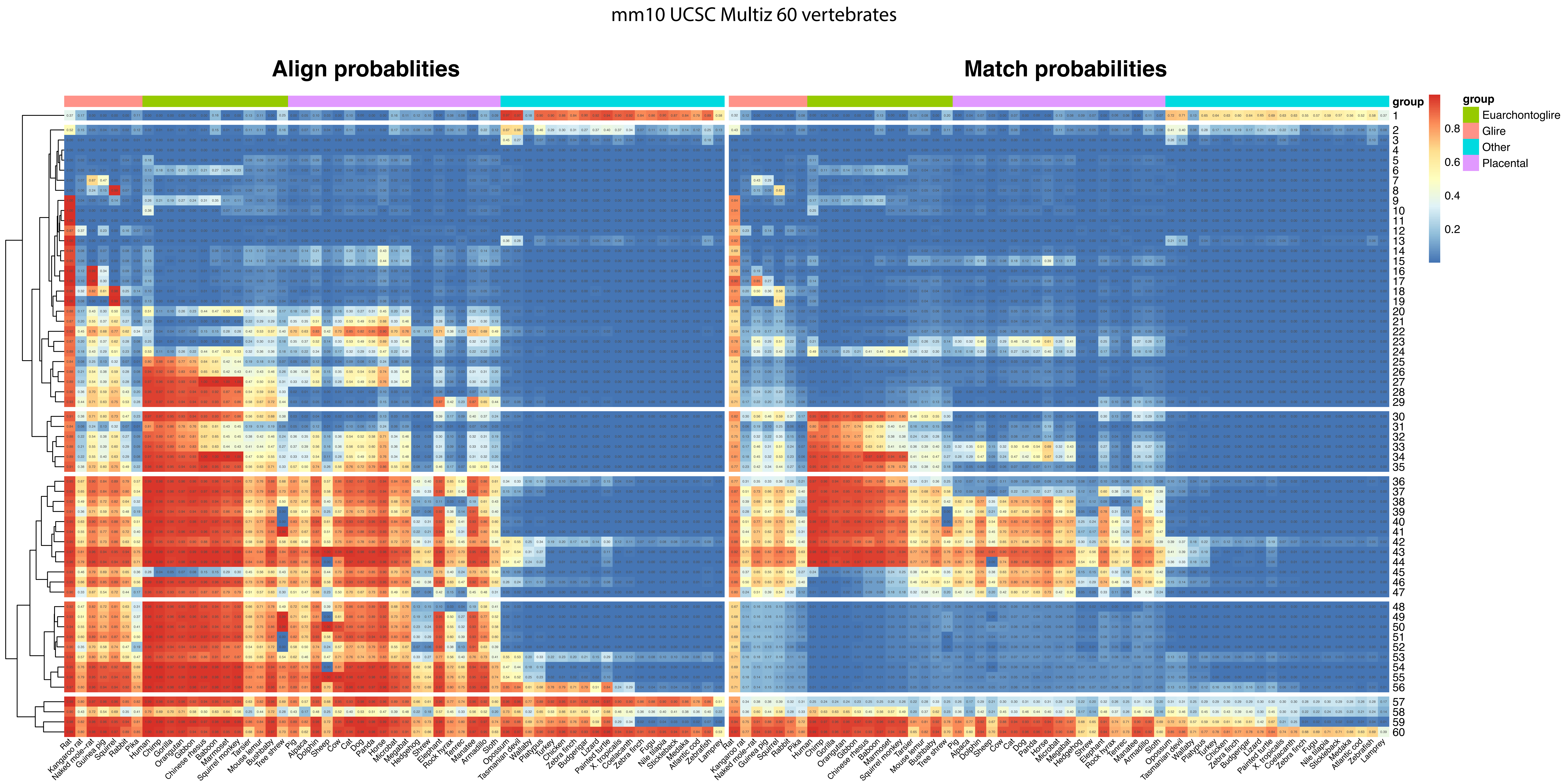

mm10 UCSC Multiz 60 vertebrates
Align probablities
Match probabilities
group
0.8
0.6
0.4
0.2
group
Euarchontoglire
Glire
Other
Placental
Rat
Kangaroo rat
Naked mole-rat
Guinea pig
Squirrel
Rabbit
Pika
Human
Chimp
Gorilla
Orangutan
Gibbon
Chinese rhesus
Baboon
Marmoset
Squirrel monkey
Tarsier
Mouse lemur
Bushbaby
Tree shrew
Pig
Alpaca
Dolphin
Sheep
Cow
Cat
Dog
Panda
Horse
Microbat
Megabat
Hedgehog
Shrew
Elephant
Rock hyrax
Tenrec
Manatee
Armadillo
Sloth
Opossum
Tasmanian devil
Wallaby
Platypus
Turkey
Chicken
Zebra finch
Budgerigar
Lizard
Painted turtle
X. tropicalis
Coelacanth
Zebra finch
Fugu
Nile tilapia
Stickleback
Medaka
Atlantic cod
Zebrafish
Lamprey

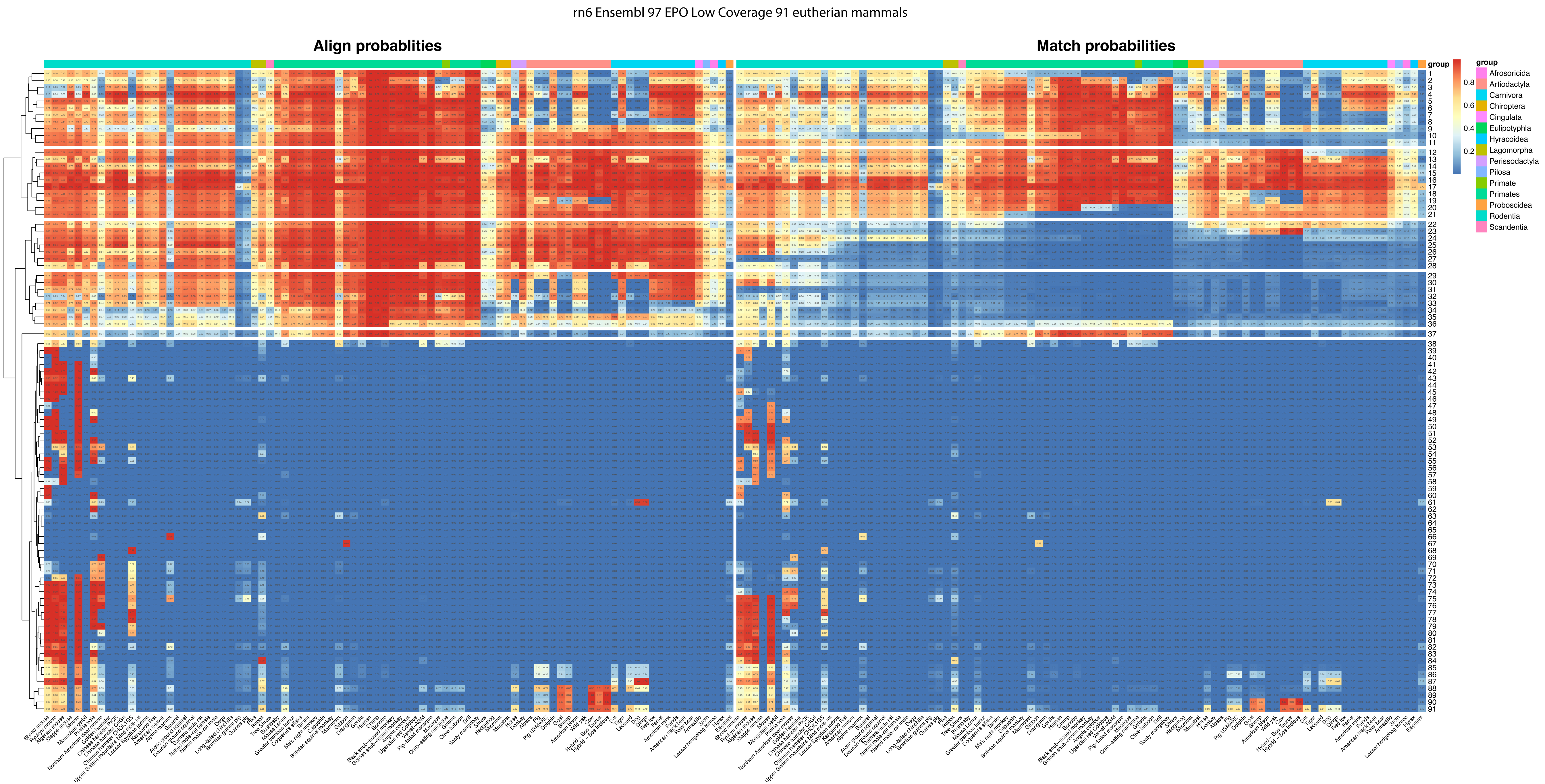
rn6 Ensembl 97 EPO Low Coverage 91 eutherian mammals
Align probablities
Match probabilities
group
0.8
0.6
0.4
0.2
group
Afrosoricida
Artiodactyla
Carnivora
Chiroptera
Cingulata
Eulipotyphla
Hyracoidea
Lagomorpha
Perissodactyla
Pilosa
Primate
Primates
Proboscidea
Rodentia
Scandentia

### rn6 Ensembl 97 EPO 38 mammals

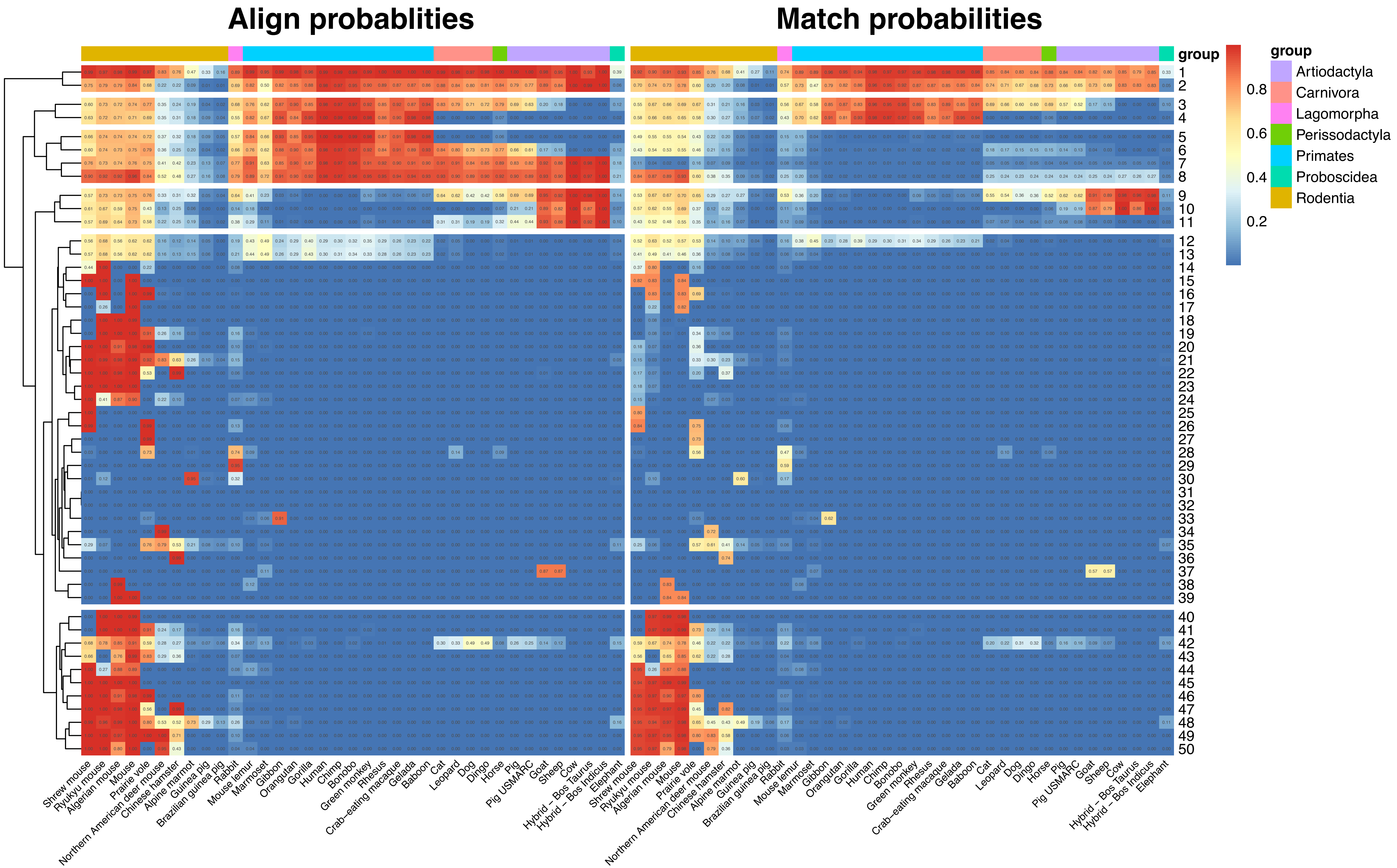

rn6 Ensembl 97 Pecan 54 amniota vertebrates

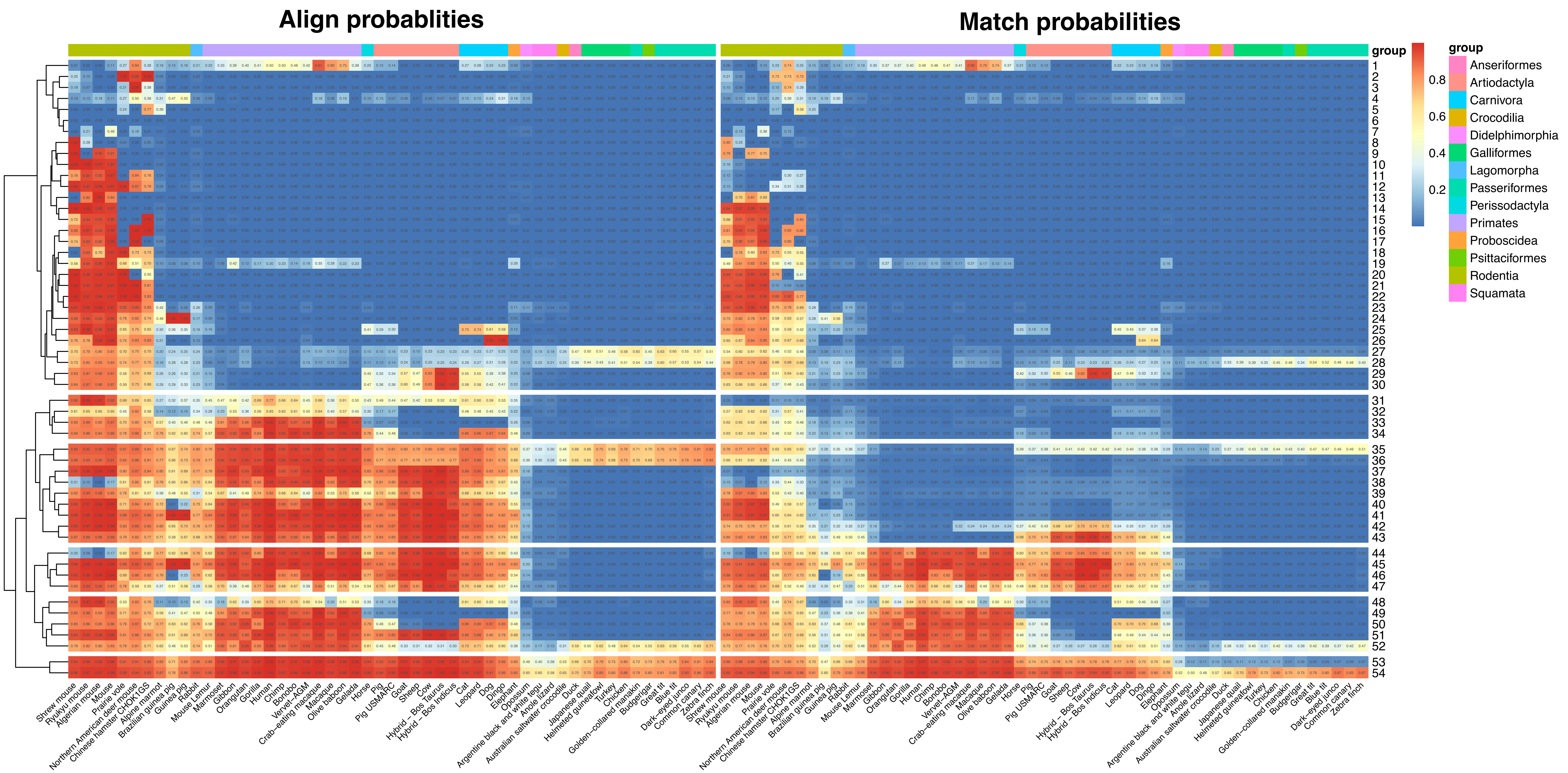

### rn6 UCSC Multiz 20 vertebrates

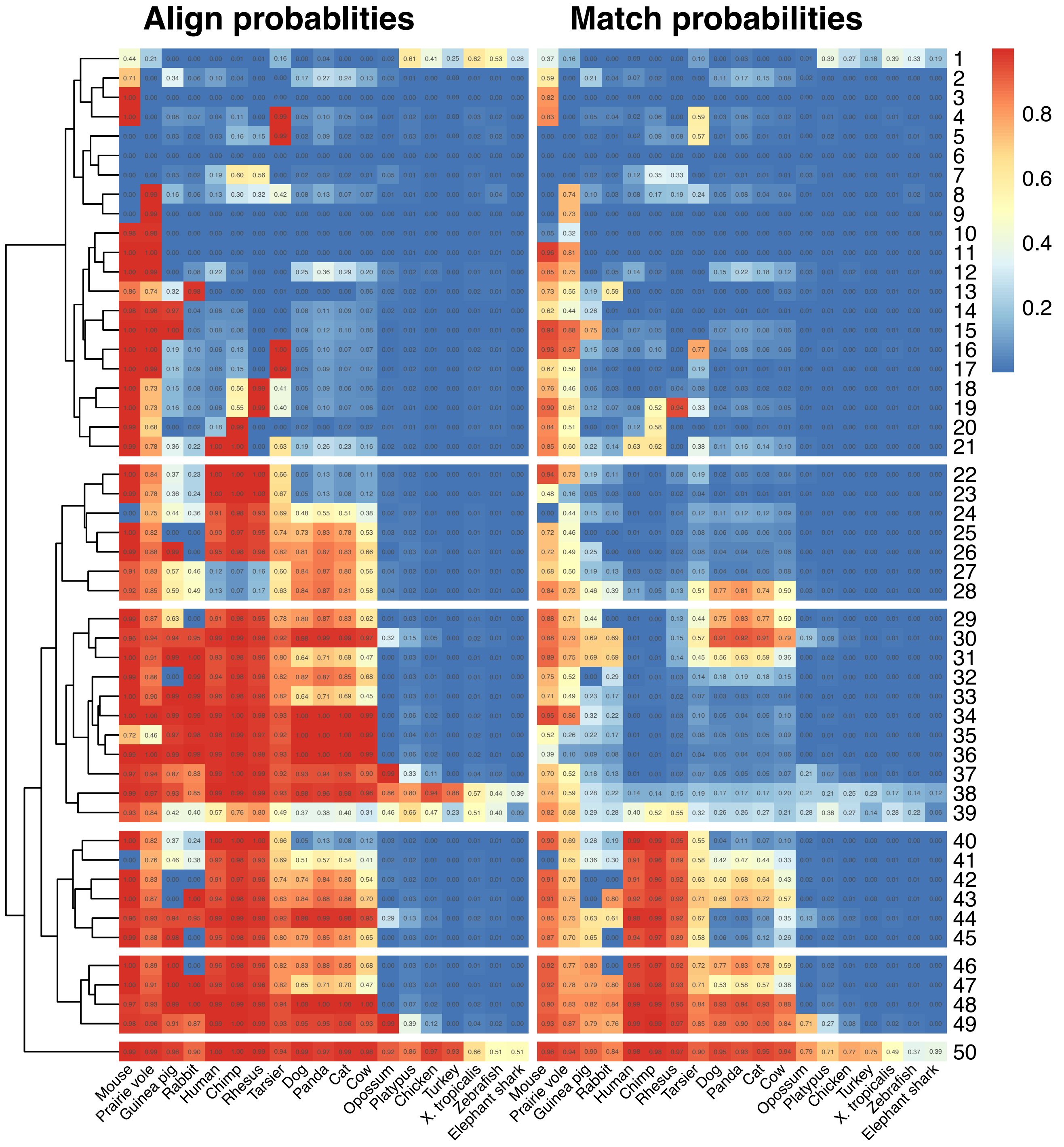

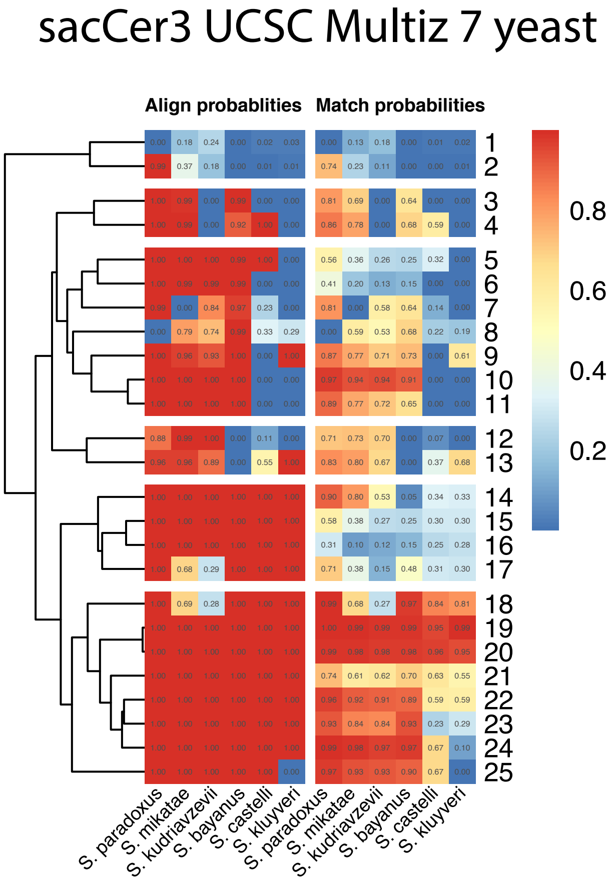
sacCer3 UCSC Multiz 7 yeast
Align probablities
Match probabilities
S. paradoxus
S. mikatae
S. kudriavzevii
S. bayanus
S. castelli
S. kluyveri
0.8
0.6
0.4
0.2
