## Supplementary Figures and Tables for "ConsHMM Atlas: conservation state annotations for major genomes and human genetic variation"

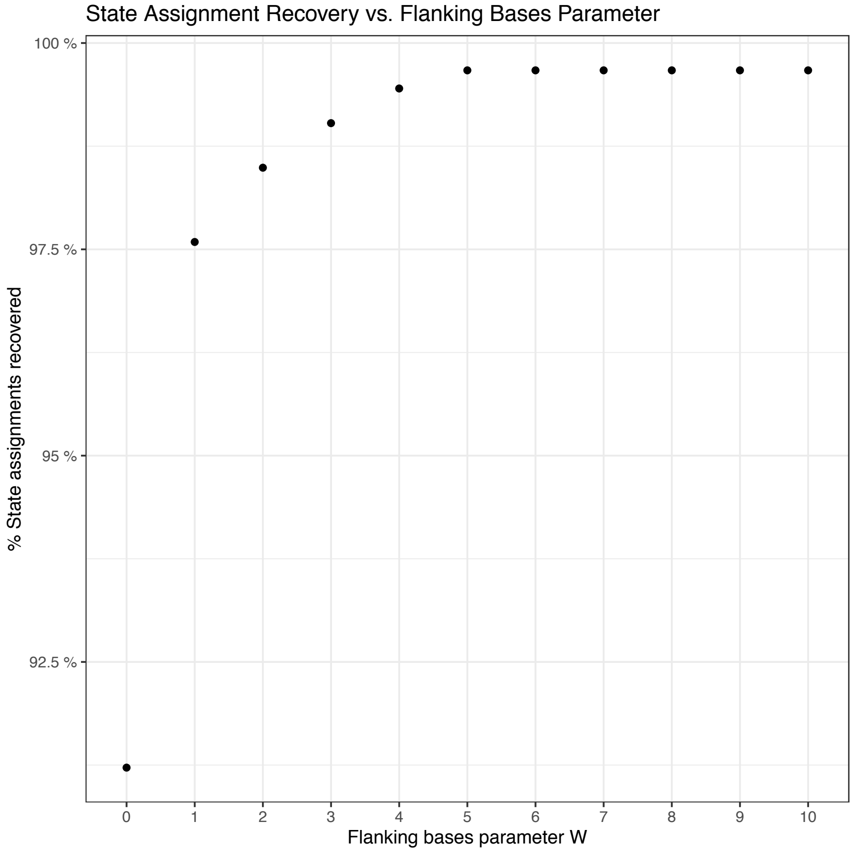


**Supplementary fig. S1: Assessment of the accuracy of allele specific conservation state assignments.** The agreement between state assignments using the segmentation of a small local window centered around a variant and the segmentation of 200kb segments. The x-axis shows the value *W*, which corresponds to *W* bases upstream and *W* bases downstream of the variant being used for the segmentation. The y-axis represents the percentage of the 40,000 variants tested for which the alternate allele gets assigned to the same state with the two approaches (**Materials and Methods**). This comparison was performed for the 100 state ConsHMM model trained on a 100-way multiple sequence alignment of 99 other species to the hg38 human genome.

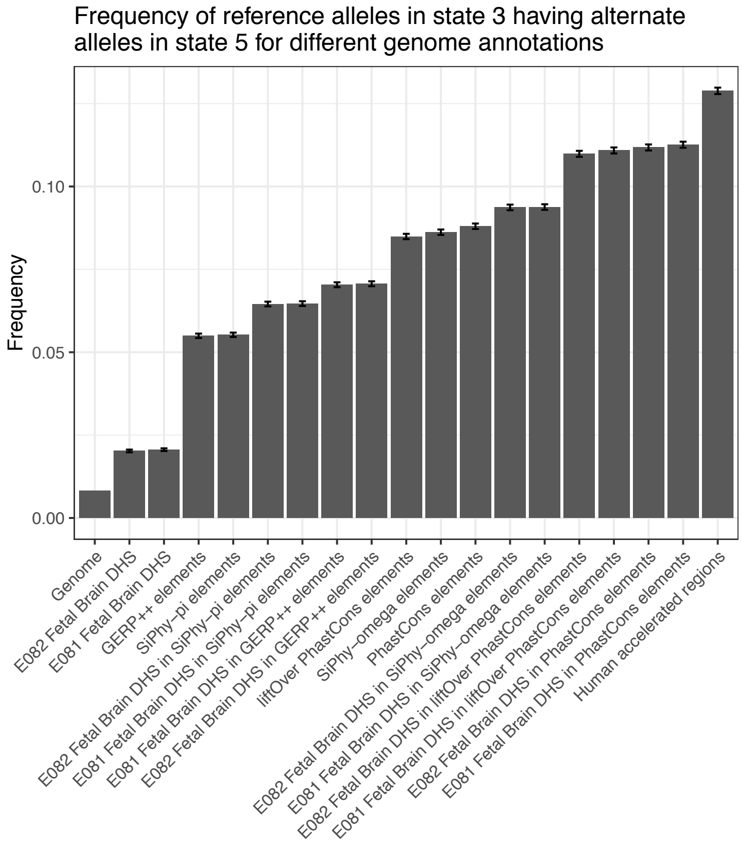


**Supplementary fig. S2: Extended example of additional information within allele specific conservation state assignments.** Shown in this plot is the frequency that state 36 assignments based on the reference allele change to state 5 over all possible alternate alleles as was done in **fig. 2**. Two sets of peaks from the Fetal Brain DNase I Hypersensitivity (DHS) assays were used (Roadmap identifiers E081 and E082) and the conserved elements used were GERP++, PhastCons, SiPhy-Pi and SiPhy-omega. Two sets of PhastCons elements were included: the elements called on the 100-way multiple sequence alignment of 99 vertebrates to the hg38 human genome, and the elements called on the 100-way multiple sequence alignment of 99 vertebrates to the hg19 human genome, which were lifted over to hg38. GERP++, SiPhy-pi and SiPhy-omega elements were all lifted over to hg38. Also shown are human accelerated regions (HAR) (Lindblad-Toh et al. 2011) and the intersection of each DHS set with each constrained element set. The HAR and DHS annotations were also lifted over to hg38.

| Organism | Target assembly | Browser | Alignment Method | Alignment |
|---|---|---|---|---|
| C. elegans | ce11 | UCSC | MultiZ | 25 nematode genomes with C. elegans |
| D. melanogaster | dm6 | UCSC | MultiZ | 26 insects with D. melanogaster |
| Dog | canFam3/CanFam3.1 | Ensembl Release 97 | EPO | 38 mammals |
| Human | Hg38/GRCh38 | UCSC | MultiZ | 99 vertebrate genomes with human |
| Human | Hg38/GRCh38 | UCSC | MultiZ | 30 mammalian (27 primate) genomes with human |
| Human | Hg19/GRCh37 | Ensembl Release 75 | PECAN | 21 amniota vertebrates |
| Human | Hg19/GRCh37 | Ensembl Release 75 | EPO_LOW_COVERAGE | 37 eutherian mammals |
| Human | Hg38/GRCh38 | Ensembl Release 97 | PECAN | 54 amniota vertebrates |
| Human | Hg38/GRCh38 | Ensembl Release 97 | EPO | 38 mammals |
| Human | Hg38/GRCh38 | Ensembl Release 97 | EPO_LOW_COVERAGE | 91 eutherian mammals |
| Mouse | mm10/GRCm38 | UCSC | MultiZ | 59 vertebrate genomes with mouse |
| Mouse | mm10/GRCm38 | Ensembl Release 97 | PECAN | 54 amniota vertebrates |
| Mouse | mm10/GRCm38 | Ensembl Release 97 | EPO | 38 mammals |
| Mouse | mm10/GRCm38 | Ensembl Release 97 | EPO_LOW_COVERAGE | 91 eutherian mammals |
| Rat | rn6/Rnor_6.0 | UCSC | MultiZ | 19 vertebrate genomes with rat |
| Rat | rn6/Rnor_6.0 | Ensembl Release 97 | PECAN | 54 amniota vertebrates |
| Rat | rn6/Rnor_6.0 | Ensembl Release 97 | EPO | 38 mammals |
| Rat | rn6/Rnor_6.0 | Ensembl Release 97 | EPO_LOW_COVERAGE | 91 eutherian mammals |
| S. cerevisiae | sacCer3 | UCSC | MultiZ | 6 yeast species with S. cerevisiae |
| Zebrafish | Zv9/DanRer7 | UCSC | MultiZ | 7 genomes with zebrafish |
| Zebrafish | Zv9/DanRer7 | Ensembl Release 75 | EPO_LOW_COVERAGE | 10 teleost fish |
| Zebrafish | GRCz11/DanRer11 | Ensembl Release 97 | EPO | 25 fish |

**Supplementary table S1: List of organisms and respective multiple sequence alignments for which ConsHMM genome annotations have been generated here.** The 'Organism' column contains the species used as the reference in the multiple sequence alignment. The 'Target assembly' column contains the genome assembly version used for each organism. The 'Browser' column contains the genome browser from where the multiple sequence alignment was downloaded. The 'Alignment method' column contains the name of the multiple sequence align-ment method used to generate the alignment. The 'Alignment' column contains a summary of the species in the multiple sequence alignment. A ConsHMM annotation based on a Multiz alignment of 99 vertebrates to the hg19 human genome is available in Arneson and Ernst (2019).
